# Affinity-matured homotypic interactions induce spectrum of PfCSP-antibody structures that influence protection from malaria infection

**DOI:** 10.1101/2022.09.20.508747

**Authors:** Gregory M. Martin, Jonathan L. Torres, Tossapol Pholcharee, David Oyen, Yevel Flores-Garcia, Grace Gibson, Re’em Moskovitz, Nathan Beutler, Diana D. Jung, Jeffrey Copps, Wen-Hsin Lee, Gonzalo Gonzalez-Paez, Daniel Emerling, Randall S. MacGill, Emily Locke, C. Richter King, Fidel Zavala, Ian A. Wilson, Andrew B. Ward

## Abstract

The generation of high-quality antibody responses to PfCSP, the primary surface antigen of *Plasmodium falciparum* sporozoites, is paramount to the development of an effective malaria vaccine. Here we present an in-depth structural and functional analysis of a panel of potent antibodies encoded by the *IGHV3-33* germline gene, which is among the most prevalent and potent antibody families induced in the anti-CSP immune response and targets the NANP repeat region. Cryo-EM reveals a remarkable spectrum of helical Fab-CSP structures stabilized by homotypic interactions between tightly packed Fabs, many of which correlate with somatic hypermutation. We demonstrate a key role of these mutated homotypic contacts for high avidity binding to CSP and in protection from *P. falciparum* malaria infection. These data emphasize the importance of anti-homotypic affinity maturation in the frequent selection of *IGHV3-33* antibodies, advance our understanding of the mechanism(s) of antibody-mediated protection, and inform next generation CSP vaccine design.

## Introduction

Vaccines are critical tools for sustainable elimination of malaria, which in 2020 was responsible for 241 million infections and 627,000 deaths worldwide (World malaria report, 2021). The pressing need for an improved vaccine is underscored by the continual emergence of resistance to antimalarial compounds by the malaria parasite, *Plasmodium falciparum* (Wicht et al., 2020). In an important milestone for global health, the first vaccine for malaria, RTS,S/AS01 (RTS,S), received recommendation for widespread use in young children living in areas of moderate to high *P. falciparum* malaria transmission by the World Health Organization (WHO) in late 2021. However, the initially robust immune response and protective efficacy conferred by RTS,S are transient, as both wane rapidly after about one year. Thus, a key challenge in malaria vaccine design is the generation of highly effective and long-lived (durable) immunity.

Many malaria vaccine candidates, like RTS,S, are based on *P. falciparum* circumsporozoite protein (PfCSP), which is the primary surface antigen of *P. falciparum* sporozoites, the stage of malaria parasites infectious to humans. The structure of PfCSP comprises three domains (Fig. 1): (1) a disordered N-terminus, which contains a heparin sulfate binding site for hepatocyte attachment; (2) a central repeat region composed of 25 to 40 major (NANP) repeats, which are interspersed by a few, N-terminal minor repeats (NVDP, NPDP); and (3) a small, structured C-terminal domain. Vaccination with whole sporozoites or full-length PfCSP generates antibodies against each domain, but the NANP repeats are immunodominant (Dame et al., 1984; Enea et al., 1984; Zavala et al., 1983). Moreover, anti-NANP monoclonal antibodies (mAbs) have been shown to confer sterile protection against malaria infection in animal models through their ability to arrest sporozoite motility in the skin and to block liver infection (Flores-Garcia et al., 2019; Foquet et al., 2014; Hollingdale et al., 1984; Hollingdale et al., 1982; Raghunandan et al., 2020; Vanderberg, 1974).

**Figure 1.**
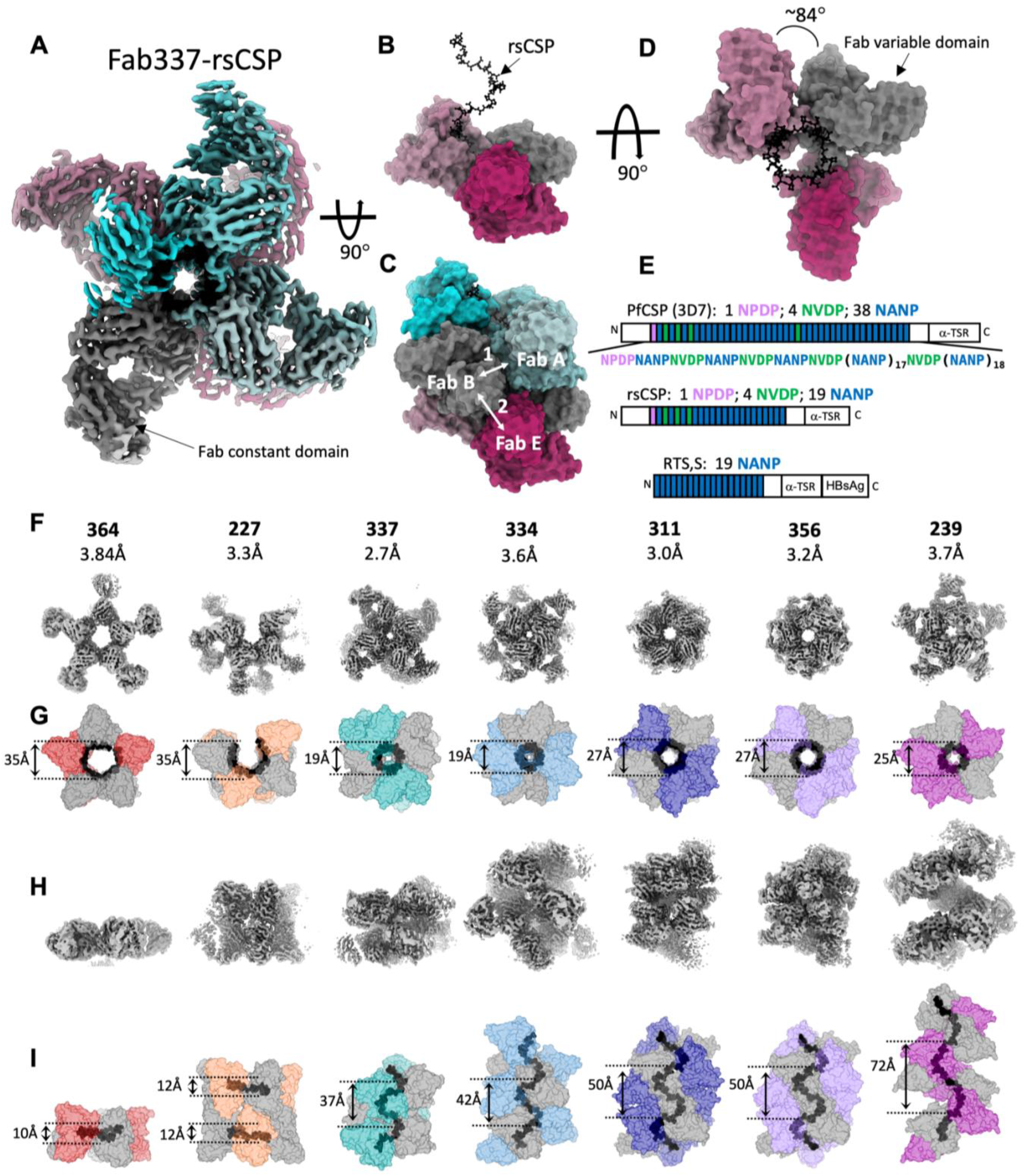
High resolution cryo-EM of *IGHV3-33* Fabs in complex with rsCSP. **(A)** Cryo-EM map of 337-rsCSP at 2.7Å, looking down the axis of the rsCSP helix. **(B)** Side-view of the 337-rsCSP structure, with four of seven Fabs removed to highlight rsCSP helical structure in black. Only variable region of Fabs are modeled. **(C)** Same as in (B), with all seven Fabs shown. Two homotypic interfaces (1 and 2) are highlighted. **(D)** Top view of (B). Rotation angle between Fabs (helical turn) is shown. **(E)** Schematic of PfCSP sequences relevant to current study. (**F)** Top view, i.e., as viewed down the axis of rsCSP helix, of cryo-EM maps. mAb name and the resolution of each cryo-EM map are listed. In panels F-I, all structures and maps are on the same scale to enable comparison of relative dimensions. **(G)** Top view of the surface representation of the various structures. rsCSP is colored in black. Diameter of the rsCSP helix is listed. **(H)** Side view of the cryo-EM maps. **(I)** Side view of various cryo-EM structures. Helical pitch is shown.

Early observation of these effects provided the rationale for the design of RTS,S (Figure 1), a virus-like particle based on the Hepatitis B surface antigen (HBsAg) that displays 19 NANP repeats and the ordered C-terminal domain of CSP (Gordon et al., 1995). Phase III clinical trials have shown that, in children aged 5-17 months, RTS,S confers modest protection (~50%) from clinical malaria at 12 months after the third vaccine dose (RTS,S CTP et al., 2011), which waned to 26% at 4 years in follow-up studies (RTS,S CTP 2015). Anti-NANP titers are associated with protection (McCall et al., 2018), and display similar induced antibody decay kinetics to other vaccines following vaccination (White et al., 2015). Thus, improving vaccine efficacy requires boosting antibody quantity over time (durability) and/or improving antibody quality (potency).

A modern approach to vaccine design entails structural analysis of potent monoclonal antibodies (mAbs) in complex with antigen (Burton, 2017). To this end, recent X-ray and cryo-EM structures have shown the repeat region is organized into NPNA structural units (Imkeller et al., 2018; Murugan et al., 2020; Oyen et al., 2017; Pholcharee et al., 2021; Pholcharee et al., 2020), and that the NPNA prolines serve as key anchor points for conserved aromatic residues in the heavy and light chain CDR loops. Interestingly, the humoral response to PfCSP is heavily biased towards antibodies descended from the human heavy chain germline gene *IGHV3-33* (Imkeller et al., 2018; Murugan et al., 2020), which has also given rise to the majority of the most potent anti-NPNA mAbs isolated to date.

We previously showed one such highly potent *IGHV3-33* mAb, mAb 311, utilizes homotypic interactions to stabilize an extended helical structure of 311 Fabs bound to rsCSP, which is a recombinant form of PfCSP containing 19 NANP repeats (Fig. 1) (Oyen et al., 2018). Somatically mutated residues mediating key homotypic contacts between adjacent Fabs were critical for stability of the extended helical structure but were not directly involved in CSP binding. Homotypic contacts were also observed in the structures of two other *potent IGHV3-33* mAbs 1210 and 239 (Imkeller et al., 2018; Pholcharee et al., 2021). Interestingly, in mAb 1210, mutations designed to disrupt these contacts significantly reduced B-cell activation in response to NANP_5_, without substantially impacting affinity to NANP_3_, implying that homotypic interactions may occur *in vivo* between adjacent B-cell receptors in response to CSP antigens. Overall, these observations suggest the nature of the NANP repeats facilitates antibody-antibody (anti-homotypic) affinity maturation, which may underlie the frequent selection of the *IGHV3-33* germline. However, whether homotypic interactions contribute to the protective efficacy of soluble antibodies, and if they occur on the surface of sporozoites, has not been demonstrated. To address these questions, we expanded our investigation of *IGHV3-33* mAbs (Pholcharee et al., 2021), and used electron microscopy combined with *in vivo* and *in vitro* assays to understand the structural basis of CSP engagement by this family of mAbs, the role of homotypic interactions, and the mechanism of protection from malaria infection.

### Helical structure formation on CSP is common among anti-NPNA mAbs from the *IGHV3-33* germline

The antibody sequences in the current study were isolated from protected individuals within the dose fractionation arm of a Phase IIa clinical trial of RTS,S (Regules et al., 2016), the same trial from which mAbs 311 and 317 were derived. We focused specifically on antibodies encoded by the heavy chain germline gene *IGHV3-33*, which has given rise to many potent anti-NPNA mAbs with a tendency toward homotypic interactions, as exemplified by Abs 1210 and 311 (Imkeller et al., 2018; Oyen et al., 2018). In all, the panel includes seven *IGHV3-33* mAbs (Table 1), including 311 for comparison, which are encoded by three different light chain genes: *IGKV1-5* (mAbs 239, 334, and 364), *IGKV3-15* (mAbs 337 and 356), and *IGLV1-40* (mAbs 311 and 227).

**Table 1.** Structural features of antibodies in this study, and helical parameters of Fab-rsCSP cryo-EM structures. Germline alleles were derived from the IMGT database, and CDR lengths are according to IMGT definitions. Full epitope describes the complete epitope of one Fab in the rsCSP cryo-EM structure. The core NPNA_2_ epitope is highlighted in green. Map res. is the overall resolution of reconstruction. The helical parameters were calculated from measurements in UCSF-Chimera. **Helical turn**: the angular step between adjacent Fabs on central rsCSP helix, as measured from the center of the helix. **Helical pitch**: the length required to complete one full helical turn, measured parallel to the rsCSP helical axis. **Helical rise:** the distance traversed along the rsCSP helix by each Fab, measured parallel to helical axis.

| mAb | IGHV Allele | IGKV/IGLV Allele | CDR3 Length (aa) |  | SHM (# aa) |  | Full epitope (cryo-EM) | Map res. (Å) | Helical Parameters |  |  |  |  |  |
| --- | --- | --- | --- | --- | --- | --- | --- | --- | --- | --- | --- | --- | --- | --- |
|  |  |  | HC | LC | HC | LC |  |  | Fabs Bound | Turn (°) | Fabs/turn | Rise (Å) | Pitch (Å) | Diameter (Å) |
| 227 | 3-33*01 | LV1-40*01 | 15 | 11 | 10 | 3 | ANPNANPNA | 3.3 | 8 | 69.9 | 5.2 | 2.3 | 12 | 35 |
| 311 | 3-33*01 | LV1-40*01 | 12 | 12 | 10 | 5 | ANPNANPNA | 3.0 | 11 | 68 | 5.3 | 10.6 | 50 | 27 |
| 239 | 3-33*03 | KV1-5*05 | 12 | 10 | 11 | 9 | NpnANPNANPNA | 3.72 | 10 | 71.9 | 5 | 14.4 | 72 | 25 |
| 334 | 3-33*08 | KV1-5*05 | 14 | 10 | 10 | 6 | NpnANPNANPNA | 3.62 | 9 | 80 | 4.5 | 9.3 | 42 | 19 |
| 364 | 3-33*03 | KV1-5*05 | 10 | 8 | 10 | 8 | ANPNANPNA | 3.84 | 5 | 61.7 | 5.8 | 1.7 | 10 | 35 |
| 337 | 3-33*08 | KV3-15*01 | 15 | 8 | 12 | 4 | ANPNANPNANP | 2.7 | 7 | 83.8 | 4.3 | 8.6 | 37 | 19 |
| 356 | 3-33*03 | KV3-15*01 | 15 | 8 | 12 | 8 | ANPNANPNANP | 3.2 | 11 | 68 | 5.3 | 10.6 | 50 | 27 |

To structurally characterize the interaction of each mAb with PfCSP, we formed complexes of the Fabs with rsCSP (Fig. 1). Initial negative-stain electron microscopy (NS-EM) imaging showed each *IGHV3-33* Fab formed well-ordered, multivalent structures on rsCSP (Fig. S1), with well-resolved Fabs radiating outwards from a central rsCSP polypeptide. For comparison, we performed the same analysis with a panel of non-*IGVH3-33*-encoded mAbs (*IGHV3-30, 3-49, 3-15*, and *1-2*) isolated from the same clinical trial (Fig. S1); we previously published EM data on two of these: 317 and 397 (Oyen et al., 2017; Pholcharee et al., 2020). Similarly to the *IGHV3-33* mAbs, the non *IGHV3-33* panel bound multivalently to rsCSP and displayed general helical or spiral curvature. However, the 2D NS-EM classes demonstrate greater structural variation and the absence of long-range helical order, in contrast to each of the *IGHV3-33* mAbs. Accordingly, we were unable to obtain stable 3D reconstructions from NS or cryo-EM of these non-*IGHV3-33* Fab-rsCSP complexes. These EM data suggest that among human mAbs, long-range helical structural ordering stabilized by homotypic interactions may be specific to the *IGHV3-33* germline.

### *IGHV3-33* antibodies exhibit a spectrum of helical conformations on rsCSP

We next utilized single particle cryo-EM to elucidate the 3D organization of these distinctive Fab-rsCSP structures, the potential roles of homotypic contacts, and the mechanisms governing their formation. Cryo-EM datasets were collected for the *seven IGHV3*-33 mAbs in our panel. Each complex was resolved to high resolution (Table S1), and the cryo-EM maps are shown in Figure 1. Of these, six structures are new, while the 311-rsCSP structure was re-refined from our previously published cryo-EM dataset to achieve higher resolution (3.0Å from 3.4Å). In general, each complex was homogeneous in both structure and composition, with the overall resolution of the reconstructions ranging from 2.7Å to 3.8Å.

This compendium of high-resolution structures reveals the remarkable conformational plasticity of PfCSP. In each complex, CSP displays some form of helical structure that is stabilized by homotypic interfaces between tightly packed Fabs bound along the length of the NPNA repeats. However, the observed helical conformations of CSP vary dramatically. These range from near planar discs with shallow pitch and large helical radius, as observed in 364 and 227, to extended helices with varying helical parameters (Table 1). Each of the extended helices in complexes with 337, 334, 311, 356, and 239 are right-handed, while the partial, disc-like helices of 364 and 227 displays left-handed curvature. The extended helical structures of CSP in the 337, 334, and 239 complexes are each unique, i.e., non-superimposable. Strikingly however, the 311 and 356 rsCSP helical structures are almost perfectly superimposable (Fig. S2), which is notable as these mAbs utilize distinct homotypic interactions and different light chains (*IGLV1-40* and *IGKV1-5*, respectively; Fig. S3). This finding suggests that either this is a relatively stable conformation of PfCSP, or that this particular structure is associated with high-level protection from malaria infection, as both 311 and 356 have been shown to be highly protective in *in vivo* mouse challenge models (Pholcharee et al., 2021).

The 227 Fab complex is distinct, as the NPNA repeats form two discontinuous, anti-parallel disc-like structures with moderate helical pitch and left-handed curvature, with each disc bound by 4 Fabs in tandem (Fig. 1). However, we note the 227 Fab structure was solved in complex with NPNA_8_ peptide instead of rsCSP, as was done for the rest of the mAbs in the panel, due to the tendency of the 227-rsCSP complex to aggregate. Thus, the two antiparallel disc structures in the 227 complex likely comprise two individual NPNA_8_ peptides, as the four available NPNA_2_ epitopes on the peptide are fully occupied by 227 Fabs and there is no density linking the two discs. Nonetheless, a NS-EM reconstruction of the 227-rsCSP complex is nearly identical to the 227-NPNA_8_ cryo-EM structure, within the 15-20Å limit of the NS data (data not shown). Therefore, this antibody may induce dimerization of separate CSP molecules mediated by homodimeric interactions of the Fabs themselves, which may have important implications for the way this antibody engages PfCSP on sporozoites.

### The *IGHV3-33* NPNA_2_ core epitope structure is highly conserved

As shown previously for this family of antibodies, the epitope of each *IGHV3-33* mAb comprises two tandem NPNA structural units, with an N-terminal type 1 β-turn followed by an Asn-mediated pseudo 3_10_ turn (Oyen et al., 2018; Oyen et al., 2017; Pholcharee et al., 2021). Interestingly, despite large differences in global helical structure, the local structure of this core (NPNA)_2_ epitope is highly conserved and exhibits a nearly identical extended S-shaped conformation in each of the seven mAbs (Fig. 2B). rsCSP binds within a deep groove running along the length of each Fab that is composed entirely of the three heavy chain CDR loops and CDRL3 (Fig. 2A). Overall, the structure of the *IGHV3-33* heavy chain is also highly conserved. Moreover, the cryo-EM structures of Fabs of 239, 356, and 364 correspond very well to our previously determined X-ray structures of these three Fabs in complex with NPNA_2_ (Fig. S2) (Pholcharee et al., 2021).

**Figure 2.**
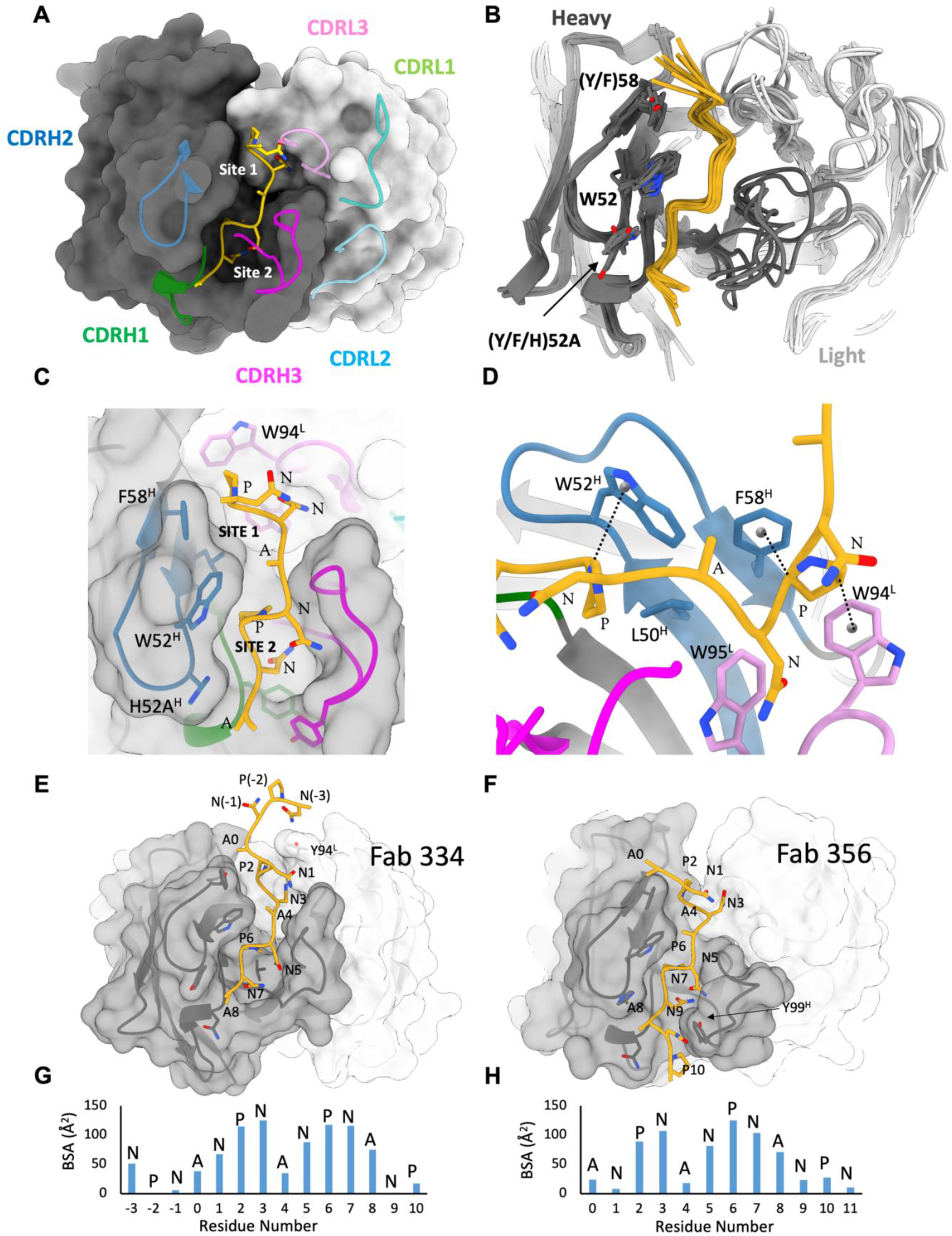
Structure and conservation of the NPNA_2_ epitope. **(A)** Surface model of a single Fab from 337-rsCSP structure, showing only the core epitope NPNA_2_ in gold. The heavy chain is colored dark gray, and light chain is in light gray. **(B)** Superposition of a Fab and NPNA_2_ from each of seven structures. Same coloring as in (A). **(C)** Zoomed-in view of paratope of 337, highlighting two hydrophobic pockets, Site 1 and Site 2. **(D)** CH-π interactions of CDRH2 and CDRL3 residues with Pro in the NPNA repeat. **(E-F)** Full epitope structure of 334 and 356, showing N and C-terminal extensions beyond NPNA_2_, which are labeled as residues 1-8. **(G-H)** Buried surface area (BSA) contributions by each residue within the full epitope of 334 (G) and 356 (H). See also Figure S2.

As noted previously for mAbs 311, 239, 356, and 364, conserved aromatic residues in CDRH2 of mAbs 227, 337, and 334 also each utilize the two prolines of the NPNA_2_ epitope as anchor points (Fig. 2B-D). The strictly conserved, germline-encoded W52 and either a Tyr (germline) or Phe at position 58 (Y/F58) each form critical, alternating CH-π interactions with the Pro of the pseudo Asn 3_10_ turn and the type 1 β-turn, respectively (Fig. 2D). These two NPNA structural units reside in two distinct hydrophobic pockets in each Fab (site 1 and site 2). While each Fab binds CSP through differing sets of interactions, this basic paratope architecture is conserved across the panel: site 1, which binds the type 1 β-turn, comprises residues from CDRH2 (Y/F58), CDRH3, and CDRL3; and site 2, which binds the pseudo Asn 3_10_ turn, is formed from the three HCDR loops and is centered on W52 and another conserved aromatic residue in CDRH2, Tyr/His52A, which in each structure packs tightly against the side chain of the C-terminal Ala of NPNA_2_ (Fig. 2C).

Importantly, the cryo-EM structures show the full epitope for a single Fab extends beyond NPNA_2_, such that adjacent Fabs engage overlapping epitopes with between 1 and 4 shared residues at the N- and C-terminal ends of each NPNA_2_ core (Fig. 2E-H; Table 1). The extent of the full epitope footprint on rsCSP tends to correlate with light chain usage and CDRH3 and CDRL3 length (Table 1). Thus, these key antibody features appear to determine the binding mode, superstructure assembly and fine epitope specificity of anti-NPNA antibodies and may also correlate with protective efficacy.

### A constellation of homotypic interactions stabilizes the CSP helical structures

Each of the multivalent antibody-CSP structures are stabilized by homotypic interactions between Fabs binding immediately adjacent NPNA_2_ epitopes, i.e., the primary homotypic interface (Interface 1; Fig. 3). This expands the full paratope, as each Fab simultaneously binds both CSP and the neighboring Fab, and substantially increases the total buried surface area (BSA) on each Fab (Table S2). The architecture of the primary homotypic interface is similar across the seven complexes and is composed mainly of the heavy chain CDR loops and CDRL3, with polar contacts between CDRH1_A_-CDRH2_B_ and CDRH3_A_-CDRL3_B_ (Fig. 3B). Importantly, this asymmetric, edge-to-edge interaction, in which FabA and FabB contribute different residues to the interface, is distinct from the asymmetric head-to-head configuration observed in the crystal structure of mAb 1210-NANP_5_, another potent *IGHV3-33/IGKV1-5* antibody (Imkeller et al., 2018). This mode of binding also differs from our previous crystal structure of 399-NPNA_6_ (*IGHV3-49/IGKV2-29*), which forms a *symmetric* head-to-head homotypic interface between adjacent Fabs (Pholcharee et al., 2021). As these latter two mAbs are not known to form stable structures on extended repeats, the edge-to-edge binding mode seen here is likely necessary for optimal geometry and packing of Fabs to promote long-range helical order.

**Figure 3.**
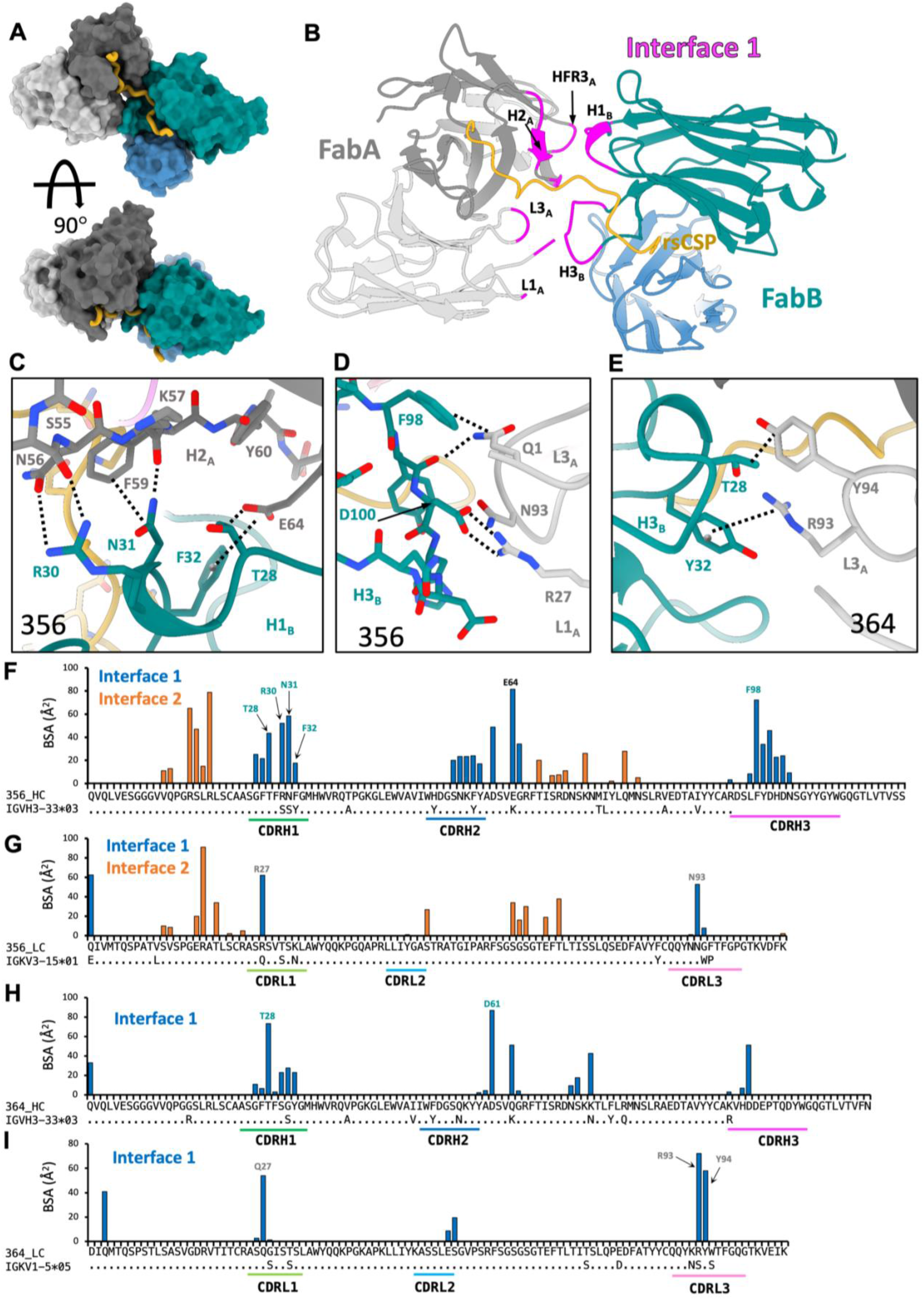
Structure of the primary homotypic interface (Interface 1). **(A)** Surface representation of two adjacent Fabs from 356-rsCSP structure. rsCSP is colored in gold. **(B)** Cartoon representation of same model as in (A), with residues mediating homotypic contacts highlighted in magenta. (**C-E)** Structural details of key homotypic interactions in 356 (C, D) and 364 (E). Specific contacts are indicated with dashed lines. **(C)** CDRH1 of FabB with CDRH2 of FabA. **(D)** CDRH3 of FabB with CDRL1 of FabA. **(E)** CDRH1 of FabB with CDRL3 of FabA. **(F-I)** Per-residue BSA contributions to homotypic interface identified in 356-rsCSP (F,G) and 364-rsCSP (H,I) structures. Note this plot does not contain BSA from CSP. See also Figures S4 and S5.

Homotypic interactions within the primary interface are derived from a diverse set of both germline-encoded residues and those that evolved through somatic hypermutation (SHM; Fig. 3F-I; Fig. S4, Table S3-S9). Two residues in CDRH1, T28 and S31, mediate key contacts between CDRH1_A_-CDRH2_B_ in nearly every complex in the panel (Fig 3C-E, Figs. S4, S5). T28 is a germline residue that is nearly strictly conserved (S28 in 337), while the S31N mutation is seen in four of the seven mAbs: 239, 311, 334, and 356 (Fig. S3). Together, these residues coordinate an extensive network of hydrophobic and electrostatic interactions, with N31 often forming multiple critical contacts with evolved basic and aromatic residues in the neighboring CDRH2_B_. Importantly, these specific interactions would likely not occur in the germline sequence (Fig. 4D-F). Other key residues in CDRH1_A_ are R30 and F32, which in both mAbs 239 and 356 form a signature motif R^30^N^31^F^32^, mutated from the germline sequence of S^30^S^31^Y^32^ (Fig. S3). In both structures, R30 forms a pair of hydrogen bonds with the N56 side chain and S55 main chain, both from CDRH2_B_, while F32 forms an anion-π bond (Philip et al., 2011) with the evolved E64 in HFR3_B_ (Fig. 3C). These interactions would also not occur upon germline reversion. Moreover, except for F32, none of the side chains of these residues directly contact CSP, providing evidence for affinity maturation to stabilize antibody-antibody rather than antibody-antigen interactions.

**Figure 4.**
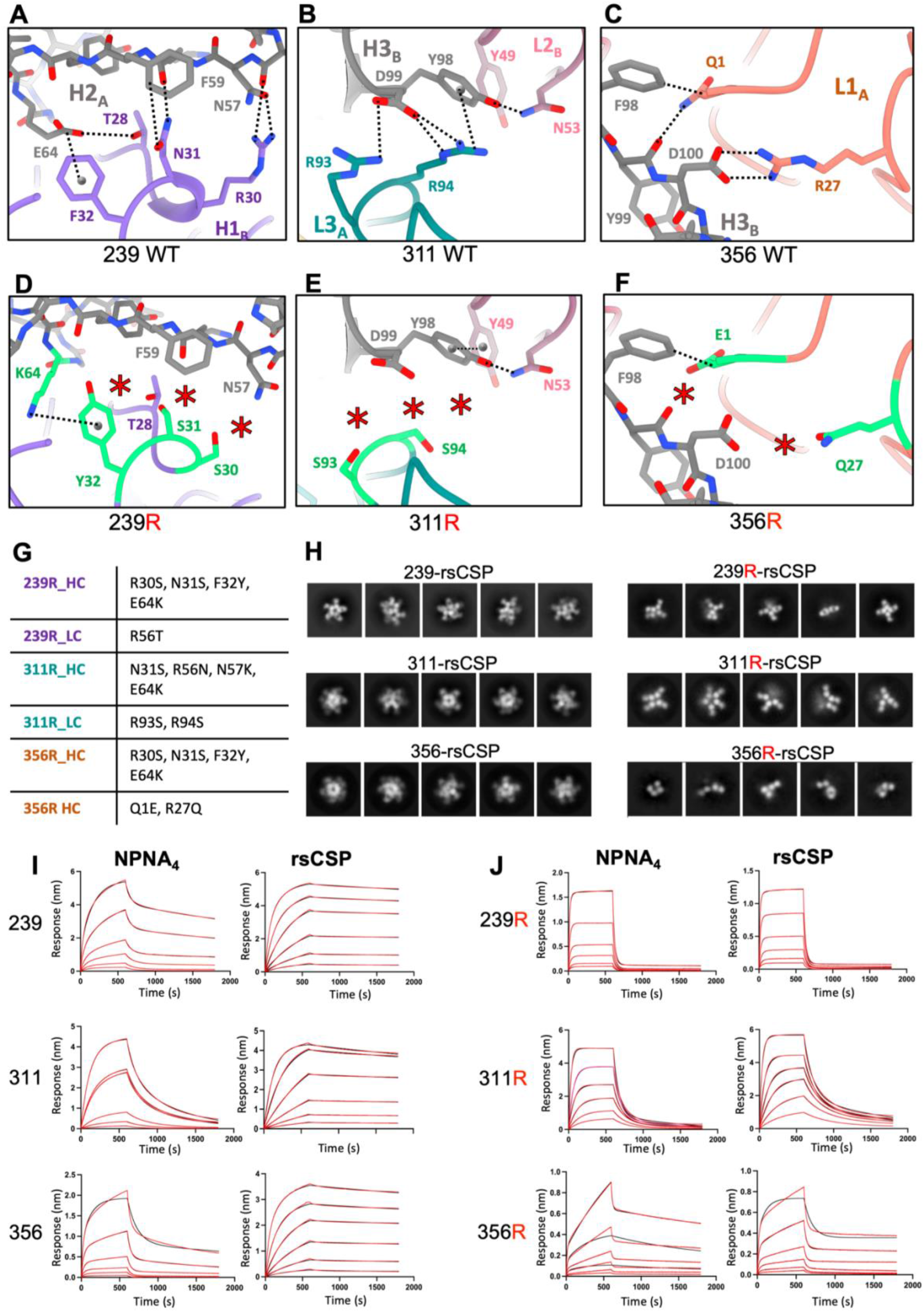
Structural and functional effects of mutagenesis of the homotypic interface. **(A-C)** Key, somatically mutated homotypic interactions observed in cryo-EM structures of 239 (A), 311 (B) and 356 (C). Dashed lines indicate observed homotypic contacts. **(D-F)** Anticipated structural impact of reversion of these residues to germline identities. Mutant structures were calculated from WT cryo-EM structures in Coot and are not experimental. Red asterisk indicates loss of homotypic contacts. Dashed lines indicate potential germline-encoded homotypic contacts. **(G)** List of germline-reverted constructs. Mutations are listed on right, using Kabat numbering system. **(H)** 2D class averages from NS-EM of WT and mutant Fab complexes with rsCSP. Mutant classes on right clearly show loss of well-ordered helical structure observed with WT Fabs. **(I,J)** Binding curves from BLI for WT (I) and mutant (J) Fabs. NPNA_4_ and rsCSP were immobilized on Streptavidin and Ni-NTA sensors, respectively, and binding of each of the Fabs were measured at 6.25, 12.5, 25, 50, 100, and 200 nM. Curves were fit with a 2:1 binding model shown in red. See also Figure S6.

As previously shown for 311, the tight helical packing in the complexes of mAbs 334, 337, and 356 creates a secondary homotypic interface between Fabs separated by about one helical turn (3 or 4 NPNA_2_ epitopes; Fig. S5). Homotypic interactions consist mostly of polar contacts within apposing heavy and light chain framework regions (FRs; Fig. S5B-D). In contrast, the secondary homotypic interface in 227 mediates 227-NPNA_8_ dimerization and defines the C2 symmetry plane for the complex (Fig. S5B). This interface is therefore symmetric and consists solely of apposing heavy chain framework residues. In general, the secondary homotypic interface contributes about half of the total BSA relative to the primary interface (Table S2). Strikingly, however, the reverse is true with 334, where the total BSA of the secondary interface is roughly twice that of the primary, suggesting a critical role for framework region residues in the stability and/or formation of this complex.

Key contacts within the secondary interface are also linked to somatic hypermutation of the germline heavy and light chain genes. In HFR1_B_ of 334, a mutated residue T19 appears critical for the interface and mediates a key hydrogen bond with S65 of LFR3_E_ (Fig S5D). In the symmetric secondary interface of 227, H82A of HFR3_B_ mediates a cation-pi bond with R75 of HFR3_G_ (Fig. S8C). Both were mutated from highly conserved residues in the germline *IGHV3-33* gene, (N82A-H and K75R). Moreover, the H82A-R75 interaction contributes nearly half of the BSA of this interface (250/550Å^2^) and defines the C2 symmetry axis of the 227-rsCSP complex (Fig. S5B,C), suggesting a critical role for this interaction. Overall, these examples represent somatic hypermutation in framework regions distal from the antigen binding site and provide further evidence for antibody-antibody affinity maturation to enhance homotypic Fab-Fab interactions.

### Mutagenesis of the homotypic interface

We have previously shown germline reversion of the somatically mutated residues that mediate homotypic contacts, but which are not directly involved in CSP binding, abrogates the 311-rsCSP helical structure (Oyen et al., 2018). To further understand the role of homotypic contacts, we applied a similar approach to two other potent mAbs in our panel, 239 and 356. Our original mutant 311 construct, 311R, has four mutations in the heavy chain (N31S, R56N, N57K, E64K), and two in the light chain (R93S, R94S). To create both 239R and 356R constructs, the RNF motif in CDRH1 was mutated to germline, along with the same E64K mutation in HFR3 (R30S, N31S, F32Y, E64K). The light chains of 239R and 356R had one and two additional mutations, respectively (239R: R56T; 356R: Q1E, R27Q) (Fig. 4G).

We first determined the impact of these mutations on binding to various CSP peptides with biolayer interferometry (BLI). We tested the hypothesis that homotypic interactions underlie the large increase in apparent affinity to peptides with increasing NANP content that is observed for this family of antibodies. Thus, we compared binding of WT and mutant Fabs to NPNA_4_, NPNA_8_, and rsCSP. In terms of NPNA_4_, the apparent affinity of 311R was essentially unchanged relative to 311 (*p*=0.17), while 356R and 239R were ~two-fold higher (improved) (*p*=,0.005) and ~two-fold lower (*p*=0.01) than 356 and 239, respectively (Fig. 4I,J; Table S10). These BLI data suggest that binding to minimal NPNA repeats is largely unperturbed by the germline mutations. As expected, for each WT Fab we observed a large increase in apparent affinity to both NPNA_8_ and rsCSP relative to NPNA_4_, largely driven by substantial reductions in the dissociation rate (k_off_). However, for the reverted mutants, both affinity and k_off_ remained roughly constant across each peptide and rsCSP. Thus, homotypic interactions are critical for high avidity binding to extended NANP repeats.

We next used NS-EM to assess the impact of the germline mutations on the structure of the Fab-rsCSP complex. As shown previously for 311R, 2D class averages of both 239R and 356R were highly variable, both in structure and stoichiometry of the Fabs (Fig. 4H). Interestingly, we observed some helical propensity in the 239R-rsCSP complex, similar to 311R, suggesting helical structure formation is at least partially germline-encoded or that CSP has a preferential bias toward a helical conformation. Nonetheless, we were unable to obtain stable 3D reconstructions for each mutant, indicative of a high degree of structural disorder. In contrast, the WT versions formed stable helical structures on rsCSP (Fig. 1, Fig. S1). Thus, somatically mutated homotypic interactions are crucial for both high avidity and for the formation and stability of long-range, helical order on rsCSP, both of which may impact protective efficacy.

To ensure these effects were due to the loss of homotypic interactions rather than unanticipated changes in the structure of the antibody paratope, which could impact the structure of the bound NPNA_2_ epitope, we solved a 1.9Å co-crystal structure of Fab311R in complex with NPNA_3_ and compared this to our previous X-ray structure of Fab311 bound to NPNA_3_ (Fig. S6) (Oyen et al., 2017). Importantly, we find the structures of both Fab and CSP peptide are nearly identical, with an overall RSMD of 0.28Å. Due to their similarity with 311R, we expect this to also be true for 239R and 356R, although we did not obtain crystals of these mAbs. Therefore, the effects of the germline mutations introduced here are likely confined to antibody-antibody binding with no significant impact on direct interactions with CSP, proving the usefulness of the germline-reverted mutants as tools to specifically probe the role of homotypic interactions.

### Affinity-matured homotypic contacts are important for high level protection

The role of homotypic contacts in protection from malaria infection is still unclear. To address this question, we compared the protective efficacy of WT and mutant 311, 239, and 356 using the liver burden assay (Fig. 5A), an *in vivo* model of malaria infection in mice that measures the ability of antibodies to prevent invasion of the liver by transgenic *P. berghei* sporozoites expressing *P. falciparum* CSP and luciferase (Flores-Garcia et al., 2019; Raghunandan et al., 2020).

**Figure 5.**
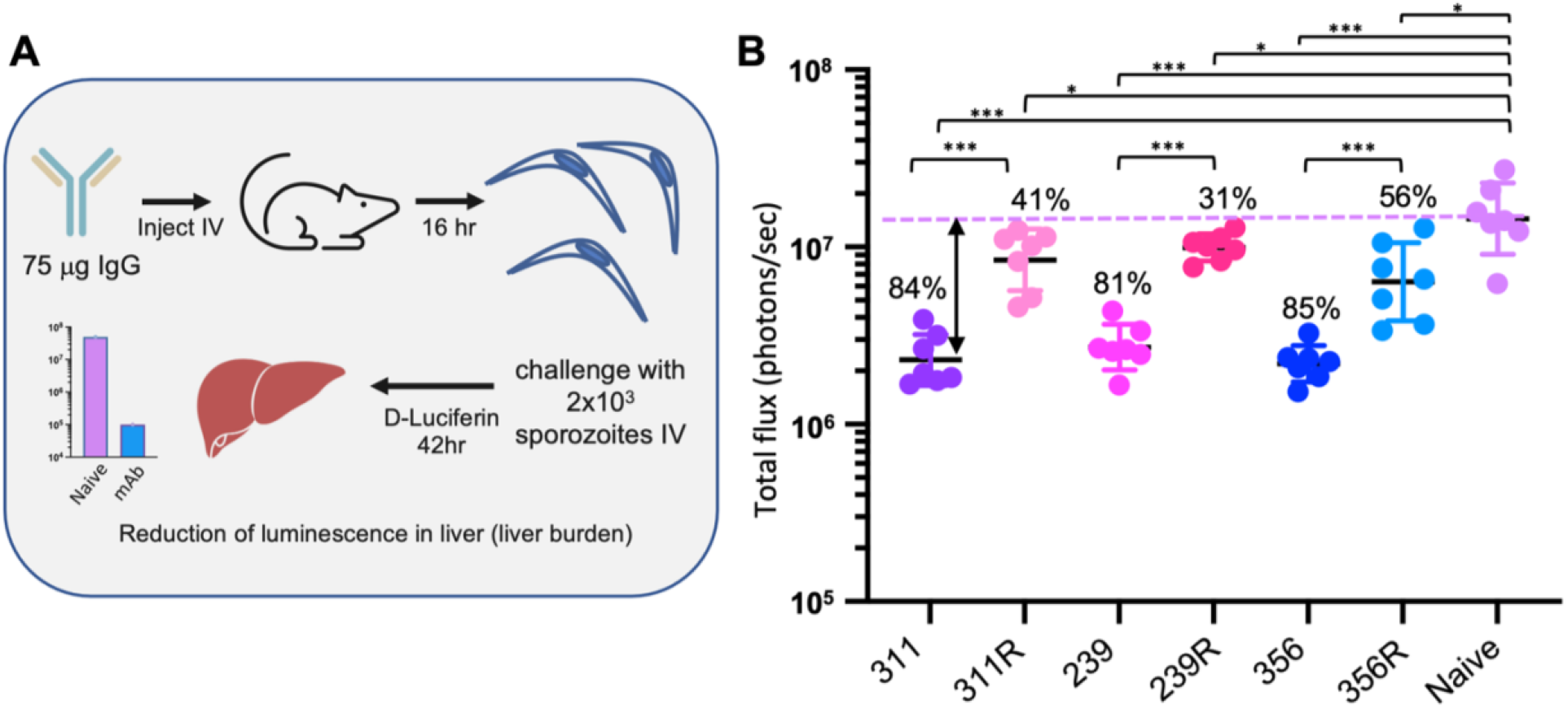
Protective efficacy of WT and germline-reverted IgGs. **(A)** Schematic of liver burden assay used to compare protective efficacy. **(B)** Liver luminescence measurements 42 hr post challenge; expressed as log total flux on the Y axis. Each group (mAb) contained seven mice. Geometric mean and SD are indicated as black and colored lines, respectively. A Mann-Whitney U-test was used to compare efficacy relative to naïve (no mAb) and between WT and mutant mAbs. Percent inhibition listed is relative to naïve. Significance: * p<0.05; *** p<0.001. See also Figure S7.

Mice were injected intravenously (IV) with 75 μg of IgG (311, 311R, 239, 239R, 356, or 356R), and 16 hours later were challenged with 2×10^3^ transgenic sporozoites. Each mAb significantly reduced parasite liver burden relative to the naïve control (Mann-Whitney U-test; *p*<0.05), which is reported as percent inhibition (Fig. 5B). Strikingly, however, 311R, 239R, and 356R each showed a consistent and dramatic reduction in percent inhibition relative to their WT counterparts, which was statistically significant in each case (Mann-Whitney U-test; *p*<0.001). In a separate experiment conducted under near identical conditions, serum IgG concentrations were measured at the time of sporozoite challenge (16hr) and were similar across the WT and variant mAbs (Fig. S7), which indicates that the differences in liver burden are likely due to differences in antibody interaction with sporozoites and not differences in antibody levels, *in vivo* mAb kinetics, or off-target responses. Overall, this is the first demonstration of a direct role of homotypic interactions in protection and implies these somatically mutated residues are critical for high-level protection from malaria, likely through their ability to mediate high avidity and helical structure formation with antibody-antibody homotypic interactions.

### Correlation of Protection and Affinity

We next compared the reduction in liver burden across each of the WT mAbs in our panel, using the same protocol as the previous protection experiment (Fig. 6A,B). For the mAbs with repeats across multiple experiments, i.e., 311, 239, and 356, and the highly-protective *IGHV3-30* mAb 317, the level of inhibition is consistent, enabling comparison of efficacy across separate experiments. As before, at 75 μg, each IgG significantly reduced parasite infectivity in the liver relative to the naïve control. While there is a range in the level of inhibition, many antibodies are highly potent and have statistically indistinguishable protection relative to mAb 317, namely mAbs 356, 311, 334, and 364. Protection for these mAbs generally ranges from around 85-92%. mAb 239 is slightly less potent than 317, at 81% inhibition, while 337 and 227 have the lowest levels of protection in the panel, at 68% and 52% inhibition, respectively (*p*<0.001; Mann-Whitney U-test). However, the reduced potency of 227 may be due to poor pharmacokinetics *in vivo* (Fig. S7). Overall, these results are consistent with our previous liver burden data testing of many of these same mAbs at 100 μg (Pholcharee et al., 2021).

**Figure 6.**
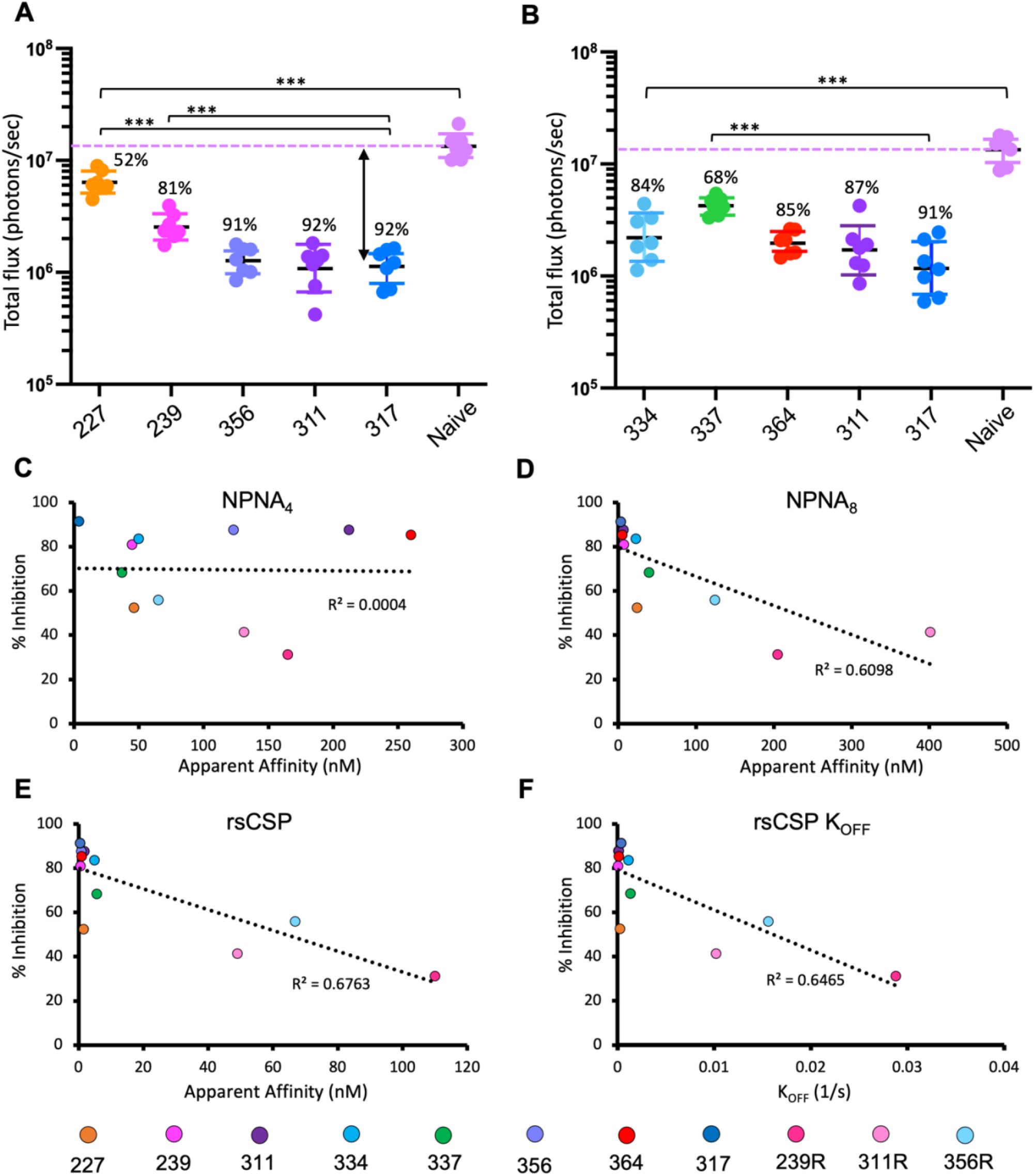
Correlation of protective efficacy and affinity to CSP. **(A,B)** Liver burden results for each mAb in the panel. Two separate experiments were conducted, and mAbs 311 and 317 were included for comparison in each. Liver burden experiments were performed and analyzed as in Figure 5B. Percent inhibition is relative to naïve (no mAb). Significance: *** p<0.001. **(C-E)** Correlation of percent inhibition with apparent affinity of each Fab, as measured by BLI, to NPNA_4_ (C), NPNA_8_ (D), and rsCSP. Binding to immobilized NPNA_4_ and rsCSP was measured at 6.25, 12.5, 25, 50, 100, and 200 nM, and binding to immobilized NPNA_8_ at 12.5, 25, 50, and 100 nM. Binding curves were fit with a 2:1 model and affinity measurements were averaged across all fits (≥4) with R^2^≥0.98. **(F)** Correlation of percent inhibition with the rate of unbinding (k_OFF_) from rsCSP in BLI experiments as in (E). See also Table S10.

As each of these mAbs target the same epitope(s) on CSP, affinity to the NPNA repeat may underlie differences in protection. To test this notion, we measured apparent affinities of each Fab to NPNA_4_, NPNA_8_, and rsCSP with BLI (Table S10). Except for the germline mutants, apparent affinity increased substantially between NPNA_4_ and NPNA_8_, and again between NPNA_8_ and rsCSP; this is likely due to the high avidity afforded by homotypic interactions, as increases in avidity were largely driven by reductions in the dissociation rate (k_off_). In general, rsCSP apparent affinity was very high (10^-9^M or higher) for the WT Fabs, and lower for the three mutants (10^-7^ to 10^-8^M). We then correlated these data with percent inhibition from the liver burden experiment. Interestingly, we observe no correlation between protection and NPNA_4_ affinity, while there is a moderate correlation with NPNA_8_ and rsCSP apparent affinity (R^2^=0.61 and 0.68, respectively), as well as rsCSP dissociation rate (R^2^=0.65). These data suggest avidity to extended NPNA repeats, which is facilitated by homotypic interactions, is a key determinant of protective efficacy among *IGHV3-33* mAbs. However, apparent affinity to NPNA_8_ or rsCSP poorly discriminates protective efficacy among the WT mAbs in the panel, which all have rsCSP apparent affinities of 10^-9^ M or lower. Therefore, high avidity to extended repeats is likely necessary but on its own insufficient to confer high level protection in *IGHV3-33* mAbs. Thus other parameters, likely concerning specifics of the interaction of antibodies with PfCSP on live sporozoites, also appear to be involved in determining protective efficacy.

## Discussion

The wealth of structural data presented herein, and the wide spectrum of observed helical conformations of rsCSP, are a vivid illustration of the extensive conformational plasticity of the NANP repeat region, which had been both predicted and demonstrated with indirect structural methods, but not directly or at high resolution (Guy et al., 2015; Kucharska et al., 2020; Patra et al., 2017; Plassmeyer et al., 2009). Our panel of cryo-EM structures reveal how these diverse conformations are anchored by a subset of key, somatically mutated residues mediating homotypic interactions across two antibody-antibody interfaces, yet which do not directly participate in CSP binding. Intriguingly, we observe this behavior in each of the seven *IGHV3-33* mAbs we examined, suggesting that, within this antibody family, affinity maturation promotes the evolution of homotypic interactions that frequently lead to long-range, ordered helical structures on CSP. Together, these data support a model in which the highly repetitive nature of the NANP repeats drives antibody-antibody affinity maturation, and that this selective advantage underlies the generation of high avidity antibodies and the frequent selection of the *IGHV3-33* germline.

We also demonstrate somatically mutated homotypic interactions, and possibly the CSP structures that they stabilize, play a key role in the mechanism of protection from *P. falciparum* infection. Specifically, we show reversion of these somatically mutated residues to their germline identities, in both heavy and light chains, abolishes well-ordered, extended helical CSP structures and eliminates the high avidity to extended NANP repeats characteristic of this antibody family, without significantly impacting affinity to the core epitope or the ability to assemble multiple Fabs onto CSP. Importantly, these effects are accompanied by a significant and consistent reduction in protective efficacy of the affinity-matured IgGs *in vivo*, relative to their WT counterparts, implying a critical role for homotypic interactions in protective efficacy for *IGHV3-33* mAbs.

Based on these data, we speculate that this family of IgGs bind multivalently on the surface of sporozoites *in vivo*, as Fabs do *in vitro*, and homotypic interactions that occur between adjacent IgGs and are critical for the stability of Ab-SPZ complexes and for protection. However, at present, little is known regarding the nature of the interaction between CSP antibodies and sporozoites, which makes it difficult to predict how differences in antibody structure or function may ultimately impact protective efficacy *in vivo*. Observations in this study and others underscore NPNA affinity alone cannot fully account for protective efficacy (Imkeller et al., 2018; Murugan et al., 2020; Pholcharee et al., 2021). Nevertheless, our functional and mutagenesis data strongly suggest high avidity to NANP repeats, driven by dramatic reductions in the off-rate, is a key component of antibody potency in the *IGHV3-33* family.

In terms of a vaccine design strategy, whether the development of homotypic interactions in an immune response is advantageous or not for both vaccine efficacy and durability remains to be determined. The current working hypothesis in the field posits that highly avid binding to extended NANP repeats, potentially afforded by homotypic interactions, induces strong B-cell activation but limits affinity maturation in germinal centers, which ultimately suppresses the development of antibodies with high affinity to the core NPNA epitope (Cockburn & Seder, 2018; Wahl & Wardemann, 2022). This would account for the robust antibody response to CSP, but also the difficulty in generating long-lived immunity and the generally low levels of somatic hypermutation observed in anti-NANP antibodies (Aye et al., 2020; McNamara et al., 2020; Murugan et al., 2018). However, there is little direct evidence showing that CSP immunogens with reduced NANP content promote the development of higher affinity antibodies, and thus improved vaccine efficacy. While two studies have indicated a trend towards greater protection in mice immunized with constructs containing reduced numbers of NANP repeats (either 9 vs 27 NANP, or 5 vs 20 NANP), the results were not statistically significant (Chatterjee et al., 2021; Langowski et al., 2020). Thus, future studies are needed to specifically assess the role of homotypic interactions in B-cell responses to CSP and whether they underlie differences in immunogenicity to different repeat constructs.

Overall, our compendium of antibody-CSP structures provides a series of high-quality structural templates which will enable both structure-based vaccine design and antibody engineering. In particular, anti-PfCSP monoclonal antibodies have emerged as promising prophylactics for malaria, with two landmark studies demonstrating the ability of two anti-PfCSP mAbs (cis43LS and L9LS) to provide months-long sterile immunity against controlled human malaria infection (CHMI) in humans (Gaudinski et al., 2021; Wu et al., 2022). Thus a key aim is to identify the most potent mAbs and improve both their potency and pharmacokinetic properties through rational, *in vitro* and *in vivo* affinity maturation. Proof-of-concept for this approach was demonstrated recently via CRISPR-based knock-in of cis43 germline heavy and light chain genes in mice (Kratochvil et al., 2021), where the authors identified a cis43 derivative with greater potency than the best-in-class mAb L9.

Our panel of cryo-EM structures will also be of immediate use in the design of NANP antigens that either promote or prevent the development of adjacent homotypic Fab-Fab interactions, or long-range bivalent IgG interactions, which may occur on extend NANP repeats. This will likely be invaluable in parsing the potentially countervailing forces homotypic interactions may exert on vaccine efficacy. Moreover, in combination with the functional and *in vivo* protection data, these structural data enable the identification of the key structural, functional, and sequence-based features of highly potent anti-PfCSP antibodies, which at this point are still not well-defined.

The fact that each of the seven *IGHV3-33* mAbs we examined, many of which are highly potent, forms distinct, extended helical structures on rsCSP suggests that these EM structures may serve as structural correlates of protection. Critically, few clear correlates of CSP vaccine-induced immunity have been identified (Julien & Wardemann, 2019; McCall et al., 2018); thus, EM-based analysis of antibody responses may be a powerful new tool for evaluating efficacy of malaria vaccines. However, at present, the correlation of higher order structures with protective efficacy is not unequivocally resolved due to the small size of our antibody panel. Future studies with larger panels of monoclonals, and especially polyclonal serum from protected and nonprotected individuals, will determine whether this phenomenon is specific to human *IGHV3-33* mAbs, or whether it represents a general solution for a productive immune response to repeat antigens.

## Acknowledgements

The authors thank B. Anderson for maintenance and administration of the cryo-EM facility at The Scripps Research Institute, and H.L. Turner and C.A. Bowman for technical support. The project was supported by National Institutes of Health grant 1F32AI150216-01A1 to G.M.M., and by funding from PATH’s Malaria Vaccine Initiative and the Bill & Melinda Gates Foundation (grant INV-004923) under collaborative agreements with The Scripps Research Institute.

This publication was also possible through support provided by the Office of Infectious Diseases, Malaria Branch, Bureau for Global Health, U.S. Agency for International Development (USAID), under the terms of Contract No. 7200AA20C00017. The opinions expressed herein are those of the author(s) and do not necessarily reflect the views of the U.S. Agency for International Development. Research by F.Z. and Y.F-G. is supported by the Bill and Melinda Gates Foundation (INV-001763), PATH’s Malaria Vaccine Initiative and the Bloomberg Philanthropies. This research used resources of the SSRL, SLAC National Accelerator Laboratory, which is supported by the U.S. Department of Energy, Office of Science, Office of Basic Energy Sciences under Contract No. DE-AC02–76SF00515. The SSRL Structural Molecular Biology Program is supported by the DOE Office of Biological and Environmental Research, and by the National Institutes of Health, National Institute of General Medical Sciences (including P41GM103393 and P30GM133894).

## Author Contributions

G.M. performed experiments, analyzed the data, prepared figures, wrote the original manuscript draft, and conceived the study. J.T., T.P., and D.O. performed experiments and analyzed the data. Y.F.G., R.M. and N.B. performed experiments, analyzed data, and wrote the manuscript. G.G., D.J., J.C., W.H.L., and G.G.P. performed experiments. D.E., R.S.M, E.L., and C.R.K. provided access to reagents and advised the study. F.Z. supervised and provided resources for the *in vivo* studies and analyzed the resulting data. I.A.W. and A.B.W. supervised the project, acquired funding, wrote the manuscript, and conceived the study. All authors contributed to manuscript editing.

## Data Availability

Cryo-EM structures and density maps were deposited to the PDB and EMDB, respectively, with the following accession codes:

**227-NPNA_8_:** 8DYT, EMD-27781

**239-rsCSP:** 8DYW, EMD-27784

**311-rsCSP:** 8DYX, EMD-27785

**334-rsCSP:** 8DYY, EMD-27786

**337-rsCSP:** 8DZ3, EMD-27787

**356-rsCSP:** 8DZ4, EMD-27788

**364-rsCSP:** 8DZ5, EMD-27789

## Materials and Methods

### CSP peptides

All peptides were produced by InnoPep Inc (San Diego, CA) at a purity level of ≥97%. Peptides for crystallography contained N-terminal acetylation and C-terminal amidation to eliminate charges at the peptide termini. Peptides for BLI were biotinylated at the C-terminus.

### Antibody sequences

All antibody sequences in the current study were derived from the MAL071 clinical trial of RTS,S/AS01 (Regules et al., 2016). Plasmablast isolation and BCR sequencing of antibody genes in malaria vaccine trials have been previously described (Regules et al., 2016; Tan et al., 2014). Fab or IgG1 heavy and light chain genes were codon-optimized and synthesized by GenScript (Piscataway, NJ).

### Protein production

Antibody genes were subcloned into pCMV or pCDNA3.4, either for expression as Fab or IgG1. Antibodies were expressed in ExpiCHO cells (Thermo Fisher) and purified using either mAb Select PrismA (GE Healthcare) or Capture Select (Thermo Fisher) columns, followed by SEC purification with a Superose S200 Increase column (GE Healthcare) equilibrated with TBS (pH 8.0). For *in vivo* testing of IgG protective efficacy in mice, endotoxins were removed with Pierce High-Capacity Endotoxin Removal Spin Columns (Thermo Fisher), following the manufacturer’s instructions. rsCSP, a recombinant, shortened construct of PfCSP containing the full N-terminal and C-terminal regions, but only 19 NANP repeats, was expressed in *E. coli* in the pET26b(+) vector, and purified as previously described (Schwenk et al., 2014).

### Mutagenesis

Inferred germline sequences were identified with IgBlast and the IMGT database. Mutations in 311R, 239R, and 356R were introduced into the light chain and Fab or IgG heavy chain by mutagenic PCR, either with the QuikChange Multi Site-Directed Mutagenesis Kit (Agilent) or the Q5 Site-Directed Mutagenesis Kit (New England BioLabs). For each point mutation, the germline codon was used. Germline reversion was confirmed by Sanger sequencing.

### Sample preparation for NS and cryo-EM

Complexes of Fabs and rsCSP, or NPNA_8_, were prepared by incubation of saturating amounts of Fab with CSP overnight at 4° C, and purified by SEC with a Superose 6 Increase column equilibrated with TBS. For negative stain EM, complexes were diluted to ~0.05 mg/mL in TBS. Sample was applied to copper grids containing a thin film of continuous carbon, made in-house, and negatively stained with 2% uranyl formate. For cryo-EM, complexes were concentrated to 2-5 mg/mL and applied to either Quantifoil holey carbon or UltrAufoil holey gold grids, and plunge-frozen with a Vitrobot MarkIV (Thermo Fisher).

### Negative stain electron microscopy

Room temperature imaging was performed either on a 120 keV Tecnai Spirit (Thermo Fisher) or a 200 keV Talos 200C (Thermo Fisher) electron microscope. Datasets on the Tecnai Spirit were collected at a nominal magnification of 52,000X (2.05Å/pix) with a Tietz TVIPS CMOS 4k x 4k camera, with a defocus of −1.5μM and a total dose of 25 e^-^/Å^2^. Datasets on the Talos were collected at a nominal magnification of 73,000X (1.98Å/pix) with a 4k × 4k CETA camera (Thermo Fisher), with a defocus of −1.5μM and a total dose of 25 e^-^/Å^2^. Leginon (Suloway et al., 2005) was used for automated data collection, and micrographs were stored in the Appion database (Lander et al., 2009). Single particle analysis was performed in RELION (Scheres, 2012), including CTF estimation, using CtfFind4 (Rohou & Grigorieff, 2015), particle picking, and reference-free 2D classification. For 3D classification, our previous negative stain reconstruction of 311-rsCSP, low-pass filtered to 60Å, was used as a reference. High quality 3D classes were used as references for 3D refinement in RELION, and C1 symmetry was used in all cases.

### Cryo-EM data collection

For 227-NPNA_8_, and 239, 334, 356, and 364 in complex with rsCSP, cryo-EM data were collected on a 200 kEV Talos Arctica (Thermo Fisher) paired with a Gatan K2 Summit direct electron detector. Micrograph movies were collected at a nominal magnification of 36,000X, resulting in a pixel size of 1.15Å, with a defocus range of −1.0 to −2.2 μm. The dose rate was ~7e^-^/pix/sec for each sample, with a total of 50 frames per micrograph movie resulting in a total dose of ~50e^-^/Å^2^. Cryo-EM data for 311 and 337 rsCSP were collected on a 300 keV Titan Krios (Thermo Fisher) with a Gatan K2 Summit direct electron detector. Cryo-EM data collection parameters for 311-rsCSP were described previously (Oyen et al., 2018), and these same data were processed in this study. For 337, imaging was performed at a nominal magnification of 29,000X (1.03Å/pix), with a defocus range of −0.9 to −2.1 μm. The dose rate was 5.3e^-^/pix/sec, and a total of 50 frames were collected resulting in a total dose of ~50e^-^/Å^2^. In all cases, Leginon was used for automated data collection.

### Single particle Cryo-EM data processing

For 311-rsCSP, our previous cryo-EM dataset was reprocessed in the current study. Raw frames were imported into RELION3.0 (Zivanov et al., 2018) and were aligned with the RELION implementation of MotionCor2 (Zheng et al., 2017). CTF estimation was performed with CtfFind4. The Laplacian-of-Gaussian picker was used for initial autopicking on a subset of micrographs, and initial 2D templates were generated with multiple rounds of 2D classification.

High quality templates were selected as input for the automated template picker in RELION for use on the whole dataset. Multiple rounds of 2D classification were used to eliminate low-quality particles, after which a total of 605,000 particles were re-extracted for 3D classification. Our previous cryo-EM reconstruction of 311-rsCSP was used as the initial reference, low-pass filtered to 60Å. A global angular search was used in the initial round of 3D classification, followed by multiple rounds of 3D classification without alignment. This process resulted in a final stack of ~400,000 particles that were re-extracted to generate a consensus refinement at 3.38Å, which is the same resolution of our previous 311-rsCSP cryo-EM map generated from these same data (EMD-9065). Further processing in RELION3.0 was used to improve the resolution of this complex. Per particle defocus values were refined in RELION, followed by another round of 3D refinement and then Bayesian polishing, which refines per-particle beam-induced motion and implements an optimized dose-weighting scheme to more accurately account for the cumulative effects of radiation damage. The resulting “shiny” particles were subjected to another round of defocus refinement and beam-tilt estimation. A final round of 3D refinement with a soft mask encompassing only the variable region of the Fabs led to the final reconstruction at 3.01Å.

A similar protocol was followed for 356-rsCSP, using the 311-rsCSP map (low-pass filtered to 60Å) as the initial model, leading to a 3.3Å reconstruction in RELION3.0. The particle stack resulting from Bayesian polishing was then imported into cryoSPARCv3.3 (Punjani et al., 2017), and two rounds of non-uniform refinement followed by global CTF (beam-tilt) refinement was performed (Punjani et al., 2020), which led to the final reconstruction at 3.2Å.

The remaining datasets were all processed according to a similar protocol in cryoSPARC. Frames were motion-corrected with MotionCor2, and the aligned and dose-weighted micrographs were imported into cryoSPARCv3.3. CTF estimation was performed with CtfFind4.

Autopicking was performed initially with the blob picker in cryoSPARC, and multiple rounds of 2D classification were used to select high quality 2D templates for subsequent template picking. Multiple rounds of 2D classification were used followed by a single round of Ab-initio reconstruction with two classes. The high-quality class was selected for further processing and was also used as the initial model. Multiple rounds of homogenous refinement, global and local CTF refinement, followed by non-uniform refinement were performed which led to the final reconstructions for each data set.

C1 symmetry was imposed for all refinements of each of the seven datasets, except for the final round of non-uniform refinement of 227, in which C2 symmetry was used. The C1 and C2 maps of 227 were nearly identical and imposing C2 improved the resolution only slightly (0.1Å).

### Model building (cryo-EM)

For Fabs 311, 239, 356, and 364, our previously-solved X-ray structures of the corresponding Fabs in complex with NPNA_2_ or NPNA_3_ were used as the starting model (PDB codes 6AXK, 6W00, 6W05, and 6WFW, respectively) (Oyen et al., 2017; Pholcharee et al., 2021). For 227, 334, and 337, an initial homology model was generated with RosettaCM (Song et al., 2013). For the heavy chain of each of these three Fabs, the heavy chain coordinates of the 311 X-ray structure (6AXK) were used as the template. To generate the light chain initial model, the light chain coordinates from the X-ray structure of the Fab with the corresponding light chain germline gene was used as the template: 311 for 227 (*IGLV1-40*), 239 (6W00) for 334 (*IGKV1-5*), and 356 (6W05) for 337 (*IGKV3-15*). The HC and LC templates were docked into the cryo-EM map, along with the NPNA_2_ peptide from 6AXK, then rebuilt and refined into the map with RosettaCM and manual adjustments with Coot (Emsley et al., 2010). Individual refined Fabs were docked into the full cryo-EM map, and the CSP peptides merged into one polypeptide chain. Further manual adjustments, if necessary, were made in Coot, and the full model was refined into the density with RosettaRelax (Conway et al., 2014).

### Structural analysis

General structural analysis, RMSD calculations, and buried surface area calculations were performed with UCSF Chimera (Pettersen et al., 2004). Homotypic contacts included in Table S3-S9 were derived from the Epitope Analyzer software, part of the ViperDB webserver (Montiel-Garcia et al., 2022). Structure figures were generated with UCSF Chimera and UCSF ChimeraX (Pettersen et al., 2021).

### 311R X-ray structure determination

311R Fab was mixed with a 5-fold molar excess of NPNA3 peptide to a final concentration of 10 mg/ml. Crystal screening was carried out using our robotic CrystalMation high-throughput system (Rigaku, Carlsbad, CA) at The Scripps Research Institute, by vapor diffusion with 0.1 μL each of protein mixture and precipitant, with 35 μL reservoir solution. 311R-NPNA_3_ crystals were grown in 0.04 M KH_2_PO_4_, 20% Glycerol, and 16% PEG3000 at 20°C and were cryoprotected in 30% glycerol. X-ray diffraction data were collected at the Stanford Synchrotron Radiation Lightsource (SSRL) beamline 12–1, and processed and scaled using the HKL-2000 package (Otwinowski & Minor, 1997) with data reduction by POINTLESS and AIMLESS (Evans, 2006). The structure was determined by molecular replacement using Phaser (McCoy et al., 2007), with the 311-NPNA3 X-ray structure (PDB 6AXK) as search model. Structure refinement was performed using Refmac5 (Kovalevskiy et al., 2018) and iterations of refinement using Coot.

### Biolayer interferometry (BLI)

BLI was performed with the Octet Red96 (ForteBio) system. A basic kinetics experiment was used to measure binding of Fabs to NPNA_4_, NPNA_8_, and rsCSP. Kinetics buffer (PBS + 0.01% BSA, 0.002% Tween-20, pH7.4) was used for all dilutions, baseline measurements, and reference subtractions. Biotinylated NPNA peptides were diluted to 5 μg/mL in kinetics buffer (KB) and immobilized onto streptavidin BLI biosensors (Sartorius); His-tagged rsCSP was diluted to ~1 μg/mL in KB and loaded onto Ni-NTA biosensors. Association and dissociation were monitored for 600 and 1200 seconds, respectively. All curves were fit with a 2:1 binding model, as there were at least two binding sites per peptide (2 sites for NPNA_4_, 4 sites for NPNA_8_, and 11 sites for rsCSP).

### Liver burden assay

The protective efficacy of IgGs in this study was assessed by the reduction in liver burden assay, as previously described (Pholcharee et al., 2021). Three separate protection experiments were conducted: one to compare the efficacy of 239R, 311R, and 356R to WT 239, 311, and 356, and two to compare efficacy of all WT mAbs in the panel to 317. Each experiment was performed under near identical conditions. Briefly, C57BL/6 mice were injected IV with 75 μg/mouse (N=7) of purified IgG and sixteen hours later challenged IV with 2000 chimeric *P. berghei* sporozoites expressing *P. falciparum* CSP and, upon liver invasion, luciferase. Forty-two hours after challenge, mice were injected with 100 μl of D-Luciferin (30 mg/mL), anesthetized with isoflurane and imaged with the IVIS spectrum to measure the bioluminescence expressed by the chimeric parasites.

### Assessment of in vivo kinetics of IgGs

Female, 6-8 week-old C57BL/6 mice were injected IV with 75μg of mAb per mouse. 16 h after injection, mice were bled and plasma was isolated. In parallel, a 384 well high binding plate (Corning 3700) was coated with anti-human IgG Fab antibody (Jackson ImmunoResearch 109-006-097) at a dilution of 1:500 and incubated overnight at 4°C. The plate was blocked with 3% BSA in PBS for 1 hr at RT. Plasma was added in a dilution series to the 384 well plate, and incubated for 1 hr at RT.Detection was measured with alkaline phosphatase-conjugated goat anti-human IgG Fcγ (Jackson ImmunoResearch 109-005- 008) at 1:2000 dilution in 1% BSA in PBS for 1hr. The plate was then washed and developed using a phosphatase substrate (Sigma-Aldrich, S0942-200TAB). Absorption was measured at 405 nm.

### Statistical Analysis

For all liver burden experiments (N=7 mice), statistical significance relative to either naïve control or between experimental conditions using the measure bioluminescence flux was assessed with a Mann-Whitney U-test, which does not assume the data can be modelled according to a probability distribution. The data were reported as the geometric mean of the total flux in the liver +/− the SD (Figure 5–6). This value was converted to percent inhibition relative to the naïve control, which is considered as 100% infected. Kinetic parameters from BLI experiments were derived from a non-linear regression of the reference-subtracted binding response according to a 2:1 binding model, as the immobilized antigen (rsCSP, NPNA_8_, or NPNA_4_) contained at least two binding sites. Values were averaged across at least 4 concentrations of Fab, and only those with R^2^>0.98 were considered (Table S10). Significance was calculated with a student’s T-test. For antibody pharmacokinetics studies in mice (N=5), non-linear regression was used to analyze the ELISA data using Prism 9 software, and circulating human IgG concentrations were interpolated based on a standard curve; data were then reported as the geometric mean +/− the SD, in mg/mL (Figure S7).

## Supplemental Figure Legends

**Figure S1.** Representative 2D class averages from negative stain EM of anti-NPNA Fabs in complex with rsCSP. To aid visualization, individual Fabs are colored representing the first (leftmost) class average of each mAb (grey).

**Figure S2.** Comparison of cryo-EM structures and previously-published Fab-peptide X-ray structures. **(A-B)** Comparison of 311-rsCSP and 356-rsCSP cryo-EM structures from this study.

**(A)** Top (left) and side (right) views of superposition of 311 (blue) and 356 (tan) complexes. **(B)** Same as in (A), showing only the rsCSP helical spiral to highlight high similarity of CSP helical structures. **(C)** Superposition of a single Fab from the 311 and 356 cryo-EM structures; CSP is in gold. **(D-G)** Comparison of cryo-EM structures of Fabs with X-ray crystal structures of corresponding Fabs bound to NPNA peptides. **(D)** 239-rsCSP cryo-EM and 239-NPNA_2_ X-ray.

**(E)** 356-rsCSP cryo-EM and 356-NPNA_2_ X-ray. **(F)**311-rsCSP cryo-EM and 311-NPNA_2_ X-ray.

**(G)** 364-rsCSP cryo-EM and 364-NPNA_2_ X-ray. **239, 356, 364:** Pholcharee et al. 2021. **311:** Oyen et al. 2017.

**Figure S3.** Multiple sequence alignment of heavy chain and light chain variable regions of the seven *IGHV3-33* mAbs in this study with inferred germline genes. **(A)** Heavy chains aligned to *IGHV3-33*01*. **(B)***IGLV1-40* light chains. **(C)** *IGKV1-5* light chains. **(D)** *IGKV3-15* light chains.

**Figure S4.** Buried surface area plots for primary and secondary homotypic interfaces for 239, 311, and 337 Fab-rsCSP cryo-EM structures. Key residues are labelled according to Kabat numbering system.

**Figure S5.** Structure of the secondary homotypic interface (Interface 2). **(A)** Surface representation of the five structures in the panel that contain a secondary homotypic interface, which exists between Fabs separated by one helical turn (Fab i and i+x). **(B)** Cartoon representation of 227 and 334, highlighting the secondary interface. rsCSP is shown as a gold surface. Only Fabs i and i+3 are shown in 334 structure for clarity. The elongated oval in 227 is the overall C2 symmetry axis of the complex. **(C)** Details of Interface 2 in 227, which is symmetric and mediated exclusively by heavy chain framework regions (FR). Local C2 axis is indicated with black oval. **(D)** Details of Interface 2 in 334 that is mediated by light chain framework region 3 (LFR3) and CDRL2 with the heavy chain framework regions 1 and 3 (HFR1 and 3). **(E-H)** BSA plots of homotypic interface 1 and 2 for 227 (E,F) and 334 (G,H).

**Figure S6.** Comparison of 311 and 311R X-ray structures. **(A)** 311-NPNA3 crystal structure (PDB 6AXK). Residues that were mutated are shown, with W52 shown for reference. CSP is in gold. **(B)** 311R-NPNA3 X-ray structure (this study). **(C)** Superposition of 311 and 311R structures. **(D)** Superposition of CSP structures from 6AXK (gray) and 311R (green). **(E)** Sequence alignment of 311 heavy and light chain variable regions with respective germline sequences. Residues mutated in 311R are shown with arrows.

**Figure S7.** Circulating concentrations of passively administered IgGs in mice, measured at time of challenge. Serum titers were measured by anti-human IgG ELISA, and calculated based on a standard curve. **(A)** WT and germline reverted 311, 239, and 356 IgG. **(B)** WT IgGs of 227, 334, 337, and 364.

**Figure S8.** Cryo-EM reconstructions of seven *IGHV3-33* mAbs in this study. **Left column**: top and side views of final EM map, colored by local resolution; color key at right. **Middle column:** representative 2D class averages. **Right column:** Fourier shell correlation for the corrected (noise-substituted) reconstruction. Note that for 311-rsCSP, cryo-EM data originally published in Oyen et al. (2018) were reprocessed here with cryoSPARCv2 and RELION3.0.

