## Supplemental Information for "Affinity-matured homotypic interactions induce spectrum of PfCSP-antibody structures that influence protection from malaria infection"

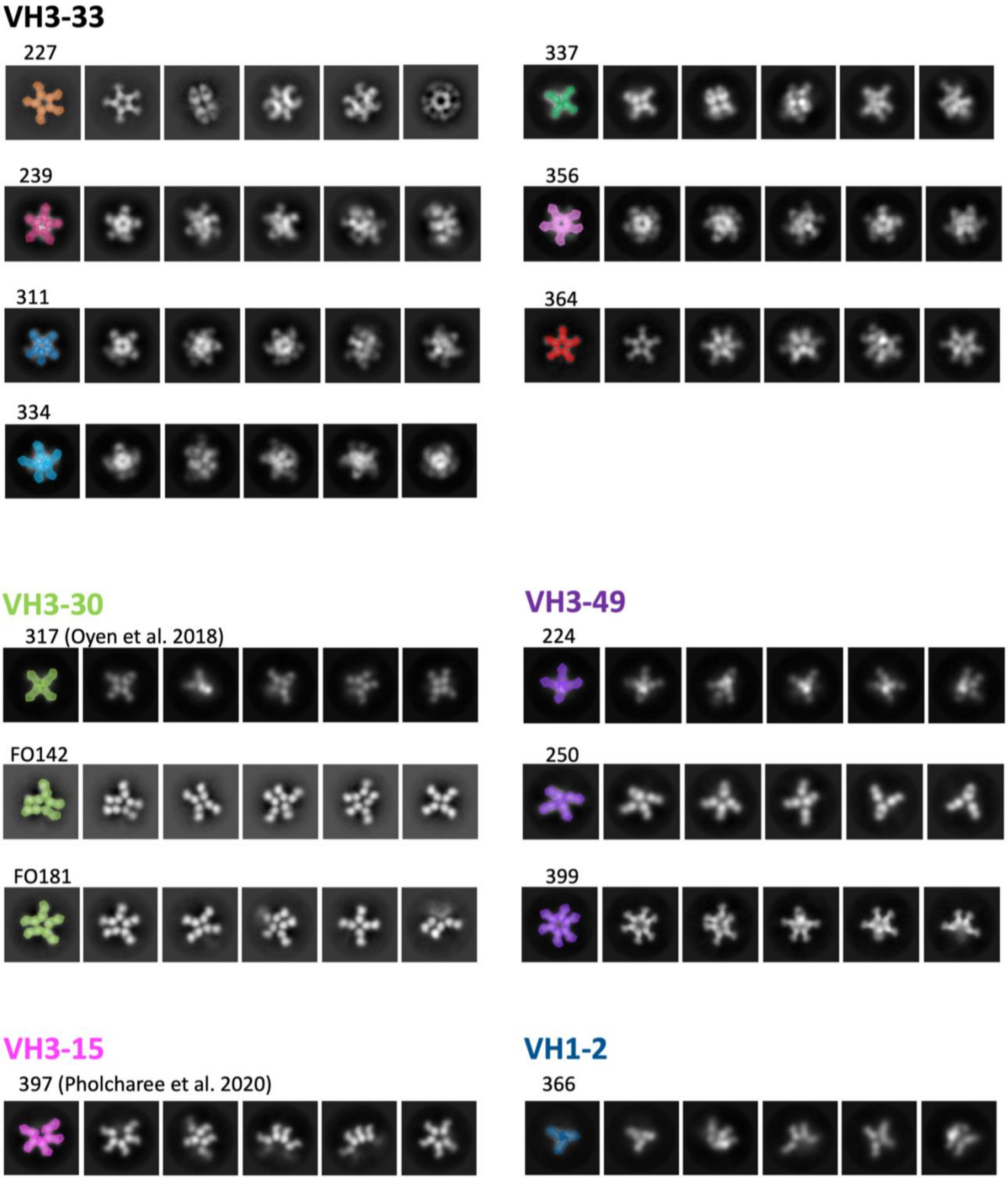

**Figure S1.** Representative 2D class averages from negative stain EM of anti-NPNA Fabs in complex with rsCSP.

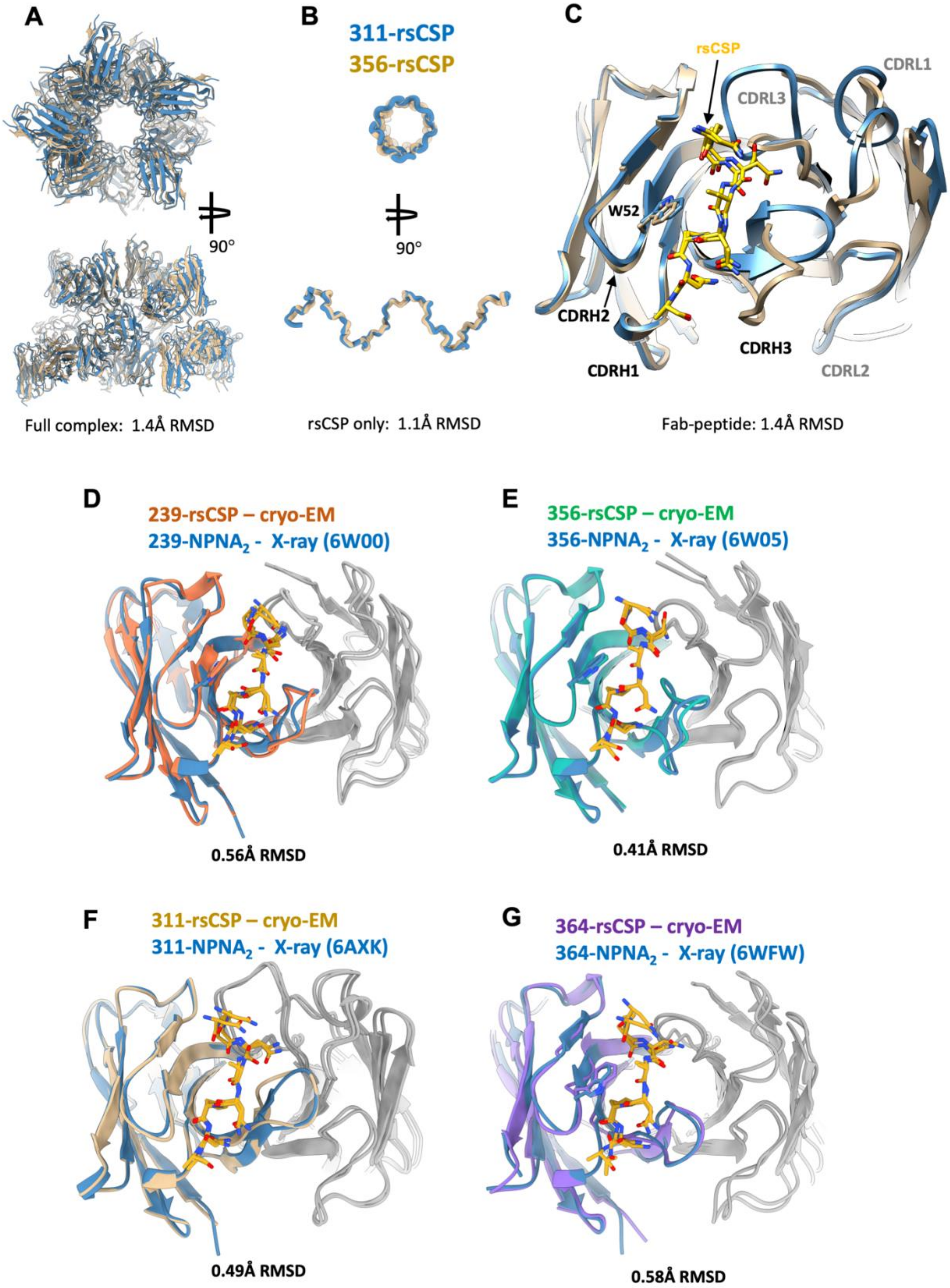

**Figure S2.** Comparison of cryo-EM structures and previously-published Fab-peptide X-ray structures.

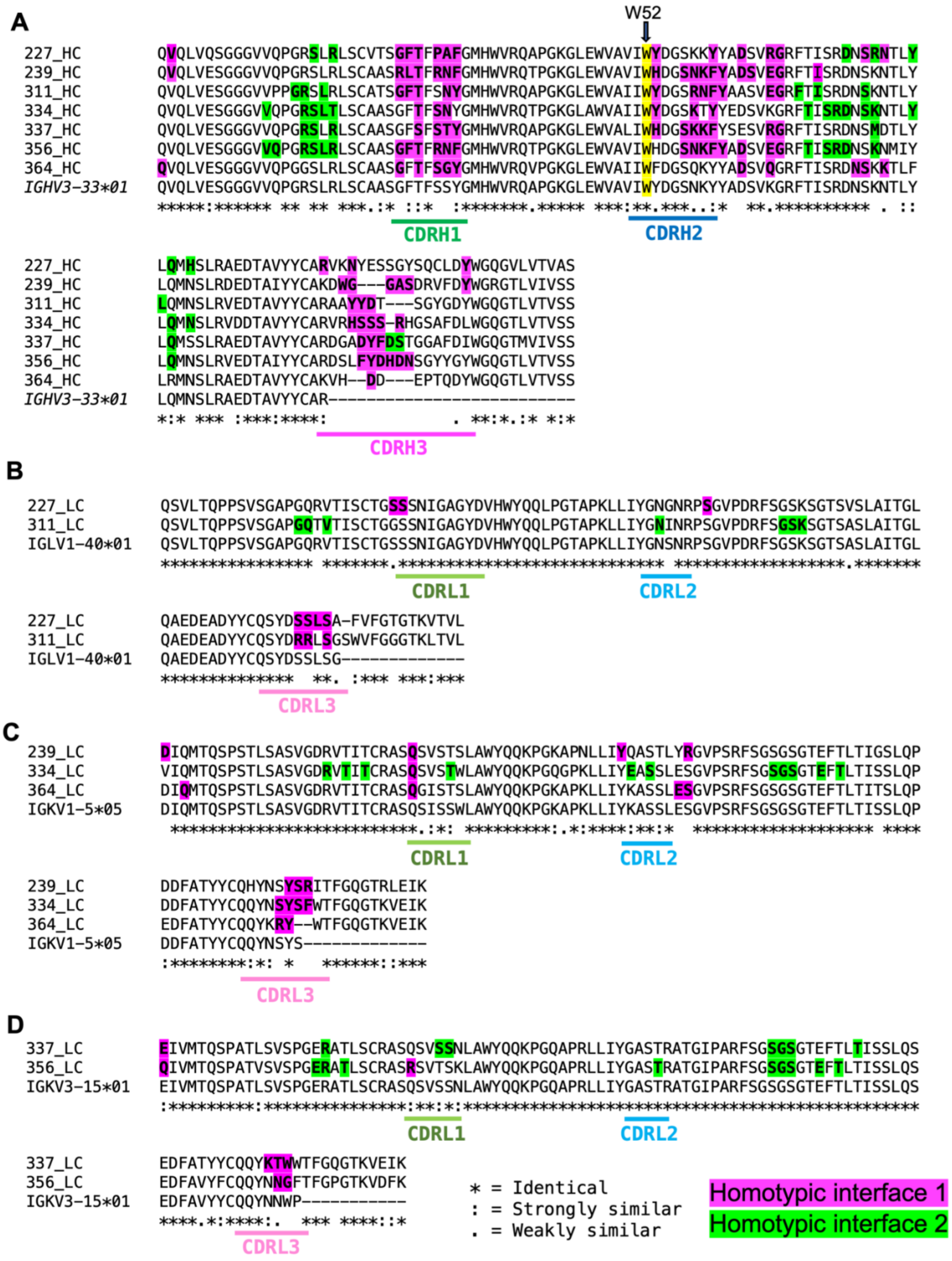

**Figure S3.** Sequence alignment of heavy chain and light chain variable regions of the seven *IGHV3-33* mAbs in this study with inferred germline genes.

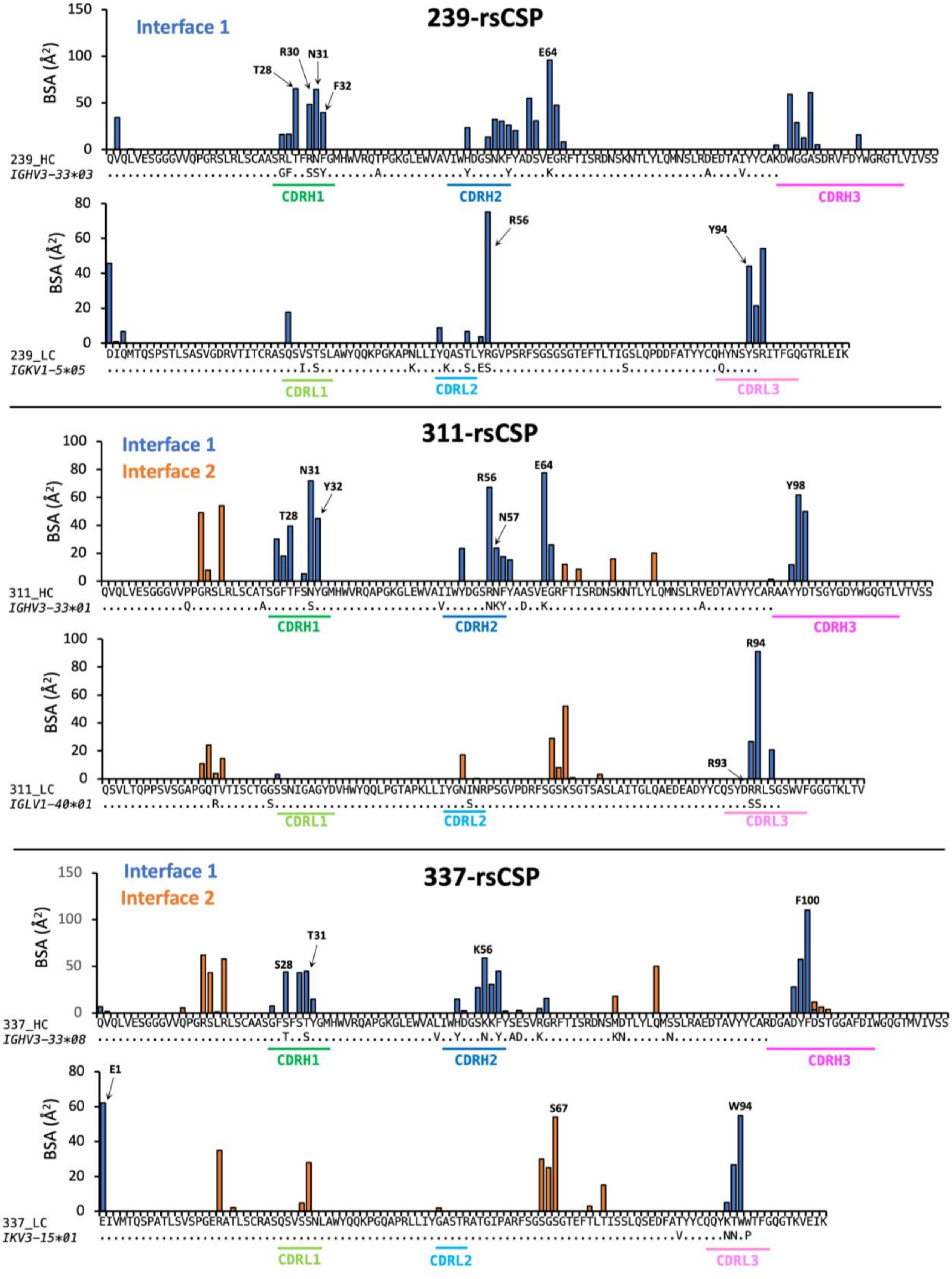

**Figure S4.** Buried surface area plots for primary and secondary homotypic interfaces for 239, 311, and 337 Fab-rsCSP cryo-EM structures.

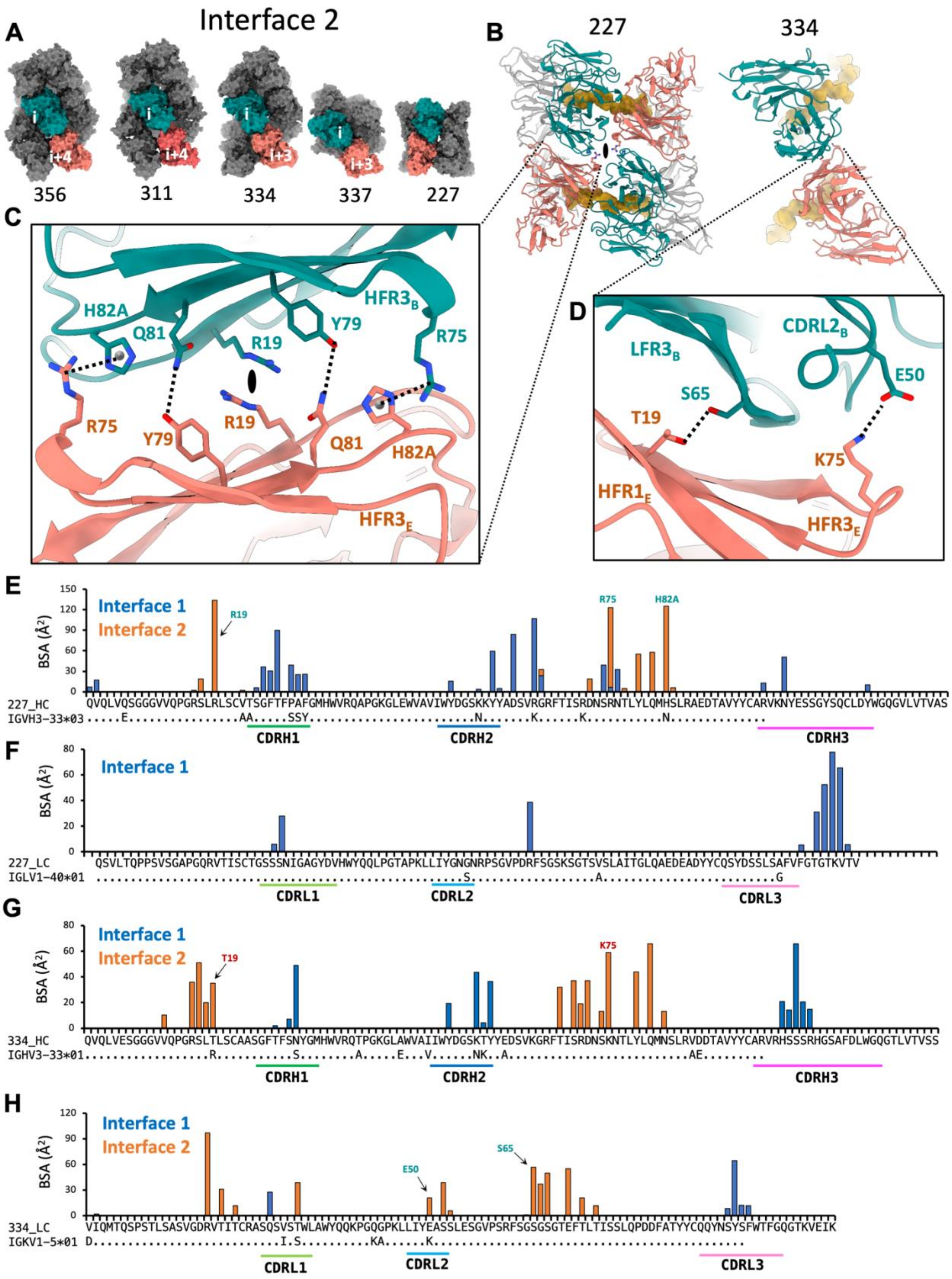

**Figure S5.** Structure of the secondary homotypic interface (Interface 2).

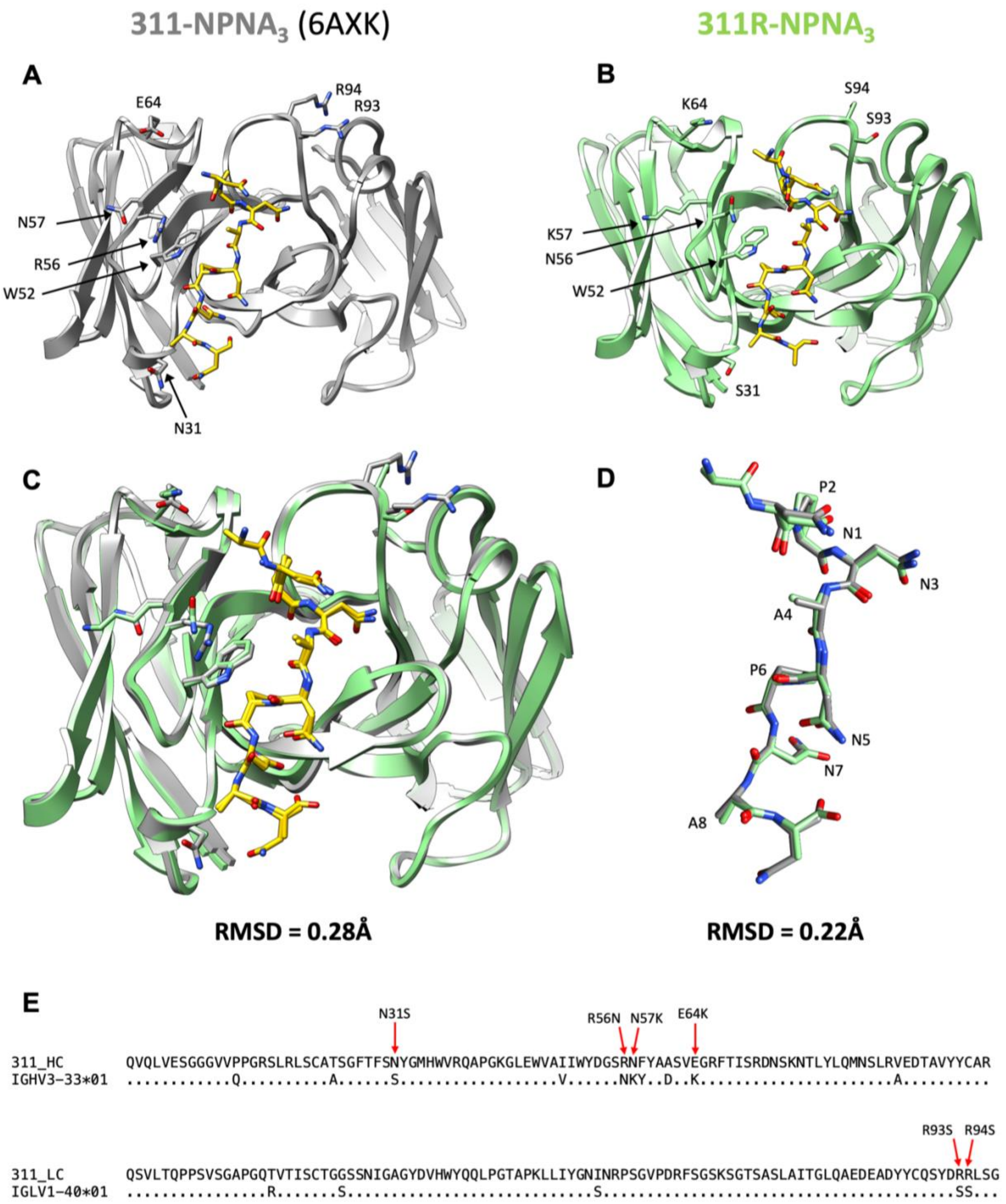

**Figure S6.** Comparison of 311 and 311R X-ray structures.

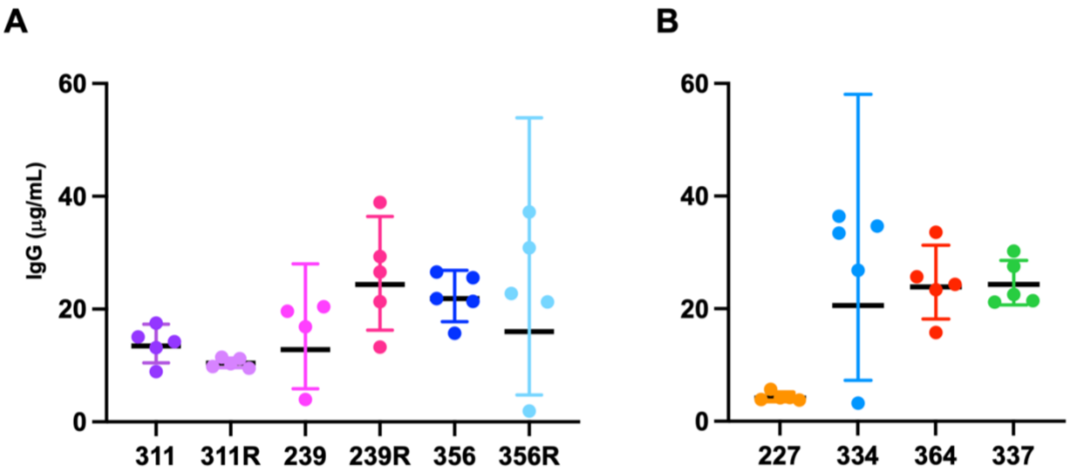

**Figure S7.** Serum titers of passively administered IgGs in mice, measured at time of challenge.

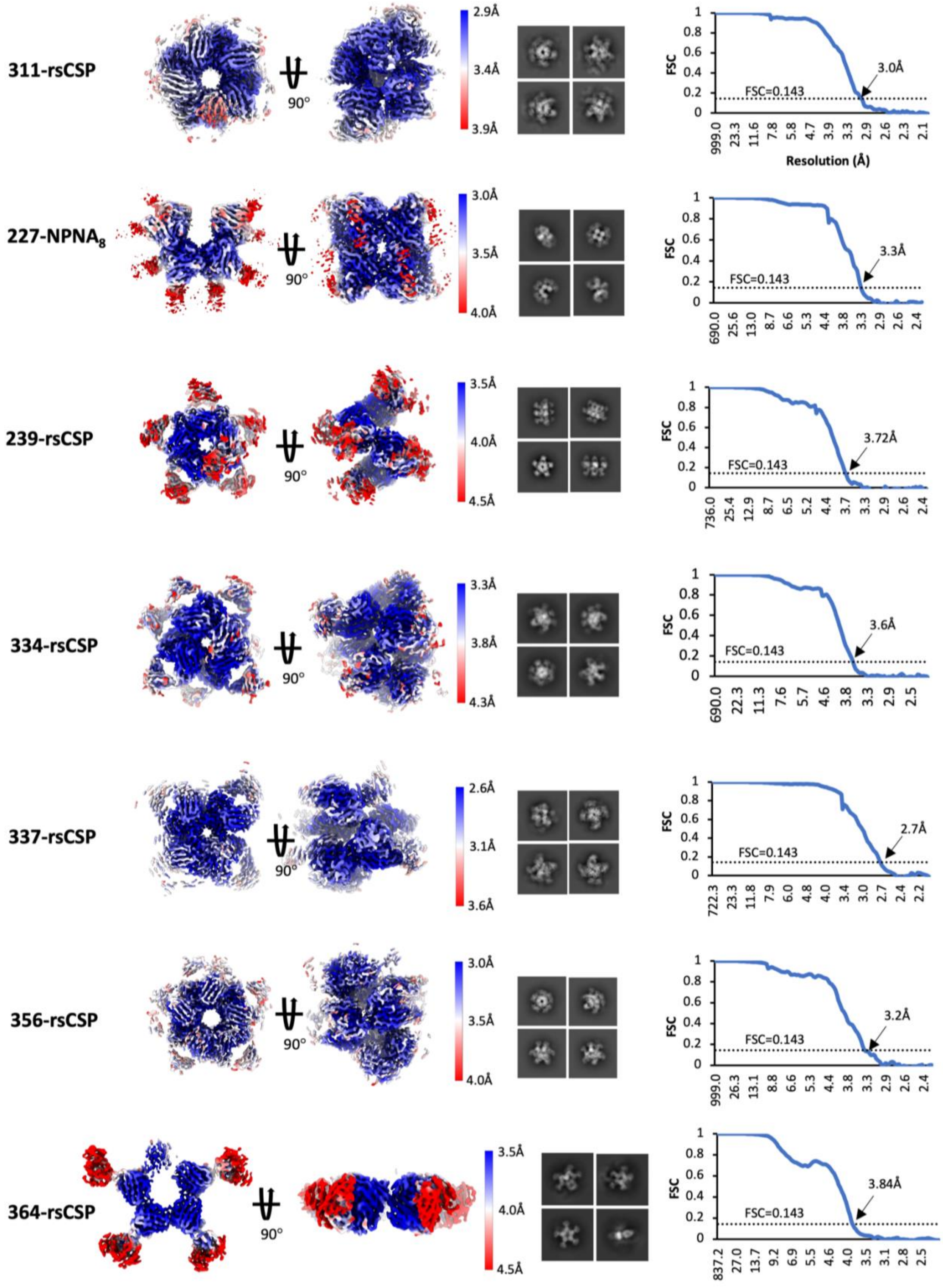

**Figure S8.** Cryo-EM reconstructions of the seven *IGHV3-33* mAbs in this study.

**Table S1.** Cryo-EM data collection parameters & model statistics

| | 227-NPNA$_8$<br>PDB: 8DYT<br>EMDB: 27781 | 239-rsCSP<br>PDB: 8DYW<br>EMDB: 27784 | 311-rsCSP<br>PDB: 8DYX<br>EMDB: 27785 | 334-rsCSP<br>PDB: 8DYY<br>EMDB: 27786 |
|---|---|---|---|---|
| **Data Collection** | | | | |
| Microscope | Talos Arctica | Talos Arctica | Titan Krios | Talos Arctica |
| Detector | Gatan K2 Summit | Gatan K2 Summit | Gatan K2 Summit | Gatan K2 Summit |
| Voltage (kV) | 200 | 200 | 300 | 200 |
| Pixel size (Å) | 1.15 | 1.15 | 1.03 | 1.15 |
| Defocus range (μm) | -0.5 to -2.0 | -1.0 to 2.2 | -0.5 to -2.5 | -1.0 to 2.2 |
| Total electron dose ($e^-/Å^2$) | 50 | 50 | 62 | 50 |
| Dose rate ($e^-/Å^2$/sec) | 5.3 | 5.3 | 5.2 | 5.3 |
| Frames per exposure | 50 | 50 | 48 | 50 |
| **Data Processing** | | | | |
| Total micrograph movies | 1595 | 1610 | 1497 | 2759 |
| Particle images in map | 70,120 | 227,439 | 399,027 | 458,706 |
| Symmetry imposed | C2 | C1 | C1 | C1 |
| Map resolution (FSC=0.143; Å) | 3.3 | 3.72 | 3.01 | 3.62 |
| Map sharpening B-factor ($Å^2$) | -122.5 | -140.6 | -54.2 | -131.5 |
| Data processing software | cryoSPARC v3.3 | cryoSPARC v3.3 | RELION3.0 | cryoSPARC v3.3 |
| **Model Refinement** | | | | |
| No. atoms in deposited model | 14,586 | 18,495 | 20,030 | 16,499 |
| RMS Deviations | | | | |
| Bond lengths (Å) | 0.007 | 0.006 | 0.003 | 0.011 |
| Bond angles (°) | 0.674 | 0.846 | 0.567 | 0.957 |
| Validation | | | | |
| Molprobity score | 1.8 | 1.93 | 1.37 | 1.92 |
| Clashscore | 8.3 | 14 | 4.74 | 10 |
| EMRinger score | 4.67 | 2.57 | 4.54 | 3.28 |
| Poor rotamers (%) | 0.45 | 0.35 | 0.33 | 0.28 |
| Ramachandran plot | | | | |
| Favored (%) | 94.9 | 95.92 | 97.35 | 94.13 |
| Allowed (%) | 5.1 | 3.65 | 2.65 | 5.87 |
| Outliers (%) | 0 | 0.43 | 0 | 0 |
| Average B-factor | 56.9 | 80.5 | 67.4 | 41.1 |

| | 337-rsCSP<br>PDB: 8DZ3<br>EMDB: 27787 | 356-rsCSP<br>PDB: 8DZ4<br>EMDB: 27788 | 364-rsCSP<br>PDB: 8DZ5<br>EMDB: 27789 |
|---|---|---|---|
| **Data Collection** | | | |
| Microscope | Titan Krios | Talos Arctica | Talos Arctica |
| Detector | Gatan K2 Summit | Gatan K2 Summit | Gatan K2 Summit |
| Voltage (kV) | 300 | 200 | 200 |
| Pixel size (Å) | 1.03 | 1.15 | 1.15 |
| Defocus range (μm) | -0.9 to -2.1 | -1.0 to -2.2 | -1.0 to -2.2 |
| Total electron dose ($e^-/Å^2$) | 50 | 50 | 50 |
| Dose rate ($e^-/Å^2$/sec) | 5.7 | 5.3 | 5.3 |
| Frames per exposure | 50 | 50 | 50 |
| **Data Processing** | | | |
| Total micrograph movies | 966 | 696 | 2049 |
| Particle images in map | 461,179 | 189,641 | 723,314 |
| Symmetry imposed | C1 | C1 | C1 |
| Map resolution (FSC=0.143; Å) | 2.68 | 3.2 | 3.84 |
| Map sharpening B-factor ($Å^2$) | -76.6 | -21.1 | -187.0 |
| Data processing software | cryoSPARC v3.3 | cryoSPARC v3.3 | cryoSPARC v3.3 |
| **Model Refinement** | | | |
| No. atoms in deposited model | 12,727 | 20,312 | 8950 |
| RMS Deviations | | | |
| Bond lengths (Å) | 0.003 | 0.002 | 0.003 |
| Bond angles (°) | 0.585 | 0.55 | 0.71 |
| Validation | | | |
| Molprobity score | 2.0 | 1.53 | 2.0 |
| Clashscore | 7.0 | 6.7 | 12.5 |
| EMRinger score | 4.1 | 4.16 | 2.21 |
| Poor rotamers (%) | 3.4 | 0.09 | 0.73 |
| Ramachandran plot | | | |
| Favored (%) | 96.5 | 97.12 | 94.6 |
| Allowed (%) | 3.5 | 2.88 | 5.4 |
| Outliers (%) | 0 | 0 | 0 |
| Average B-factor | 55.1 | 58.4 | 69.2 |

**Table S1.** Cryo-EM data collection parameters and model statistics.

Table S2. Buried surface area (BSA) calculations

| | Full Epitope BSA (Å²) | | Homotypic Interface BSA (Å²) | |
|---|---|---|---|---|
| **mAb** | **Fab BSA** | **CSP BSA** | **Interface 1** | **Interface 2** |
| 227 | 837 | 911 | 1042 | 552 |
| 239 | 964 | 1044 | 1139 | |
| 311 | 510 | 662 | 727 | 331 |
| 334 | 1058 | 1127 | 425 | 950 |
| 337 | 941 | 1110 | 714 | 455 |
| 356 | 888 | 975 | 899 | 674 |
| 364 | 878 | 955 | 738 | |

**Table S2.** Buried surface areas (BSA) for individual Fabs bound to rsCSP, and for the two homotypic interfaces in the rsCSP complex.

**Table S3.** Homotypic contacts in 227-NPNA$_8$ cryo-EM structure

| Interface | Chain Fab 1 | Residue 1 | Position 1 | Atom | Chain Fab 2 | Residue 2 | Position 2 | Distance (Å) | Predicted interaction |
|---|---|---|---|---|---|---|---|---|---|
| 1 | H | GLN | 1 | NE2-CD2 | D | LEU | 95 | 4.1 | Van-der-Waals |
| 1 | H | SER | 25 | C-OD2 | C | ASP | 61 | 3.3 | Van-der-Waals |
| 1 | H | GLY | 26 | N-OD2 | C | ASP | 61 | 2.4 | H-bond |
| 1 | H | GLY | 26 | N-OD1 | C | ASP | 61 | 4.4 | H-bond |
| 1 | H | GLY | 26 | O-CG | D | LEU | 95 | 3.6 | Van-der-Waals |
| 1 | H | PHE | 27 | N-OD2 | C | ASP | 61 | 3.7 | H-bond |
| 1 | H | PHE | 27 | O-NH1 | C | ARG | 64 | 3.6 | H-bond |
| 1 | H | PHE | 27 | CB-O | D | SER | 94 | 4.3 | Van-der-Waals |
| 1 | H | PHE | 27 | CB-O | D | LEU | 95 | 3.6 | Van-der-Waals |
| 1 | H | PHE | 27 | CB-CA | D | LEU | 95 | 4.1 | Hydrophobic |
| 1 | H | THR | 28 | OG1-CZ3 | C | TRP | 47 | 4.5 | Van-der-Waals |
| 1 | H | THR | 28 | CG2-CD1 | C | TYR | 58 | 4.5 | Van-der-Waals |
| 1 | H | THR | 28 | OG1-O | C | TYR | 59 | 4.0 | H-bond |
| 1 | H | THR | 28 | CG2-NH2 | C | ARG | 64 | 3.2 | Van-der-Waals |
| 1 | H | THR | 28 | OG1-NH2 | C | ARG | 64 | 4.2 | H-bond |
| 1 | H | THR | 28 | N-O | D | LEU | 95 | 3.1 | H-bond |
| 1 | H | THR | 28 | OG1-O | D | LEU | 95 | 3.5 | H-bond |
| 1 | H | THR | 28 | OG1-O | D | SER | 95A | 2.8 | H-bond |
| 1 | H | THR | 28 | OG1-CB | D | ALA | 95B | 3.7 | Van-der-Waals |
| 1 | H | THR | 28 | OG1-N | D | ALA | 95B | 4.3 | H-bond |
| 1 | H | PHE | 29 | N-NH1 | C | ARG | 64 | 3.9 | Van-der-Waals |
| 1 | H | PRO | 30 | CD-NE | C | ARG | 64 | 3.3 | Van-der-Waals |
| 1 | H | ALA | 31 | CB-OG | D | SER | 95A | 3.7 | Van-der-Waals |
| 1 | H | PHE | 32 | CZ-O | D | SER | 93 | 3.8 | Van-der-Waals |
| 1 | H | PHE | 32 | CZ-O | D | SER | 94 | 3.4 | Van-der-Waals |
| 1 | H | PHE | 32 | CE2-O | D | LEU | 95 | 3.9 | Van-der-Waals |
| 1 | H | PHE | 32 | CE2-CB | D | SER | 95A | 3.5 | Van-der-Waals |
| 1 | H | ASN | 73 | O-CD | C | ARG | 64 | 3.9 | Van-der-Waals |
| 1 | H | SER | 74 | O-CG | C | ARG | 64 | 3.8 | Van-der-Waals |
| 1 | H | SER | 74 | O-CA | C | GLY | 65 | 3.3 | Van-der-Waals |
| 1 | H | SER | 74 | O-N | C | GLY | 65 | 3.7 | H-bond |
| 1 | H | ASN | 76 | ND2-OD1 | C | ASP | 61 | 3.5 | H-bond |
| 1 | H | ASN | 76 | OD1-CD | C | ARG | 64 | 3.2 | Van-der-Waals |
| 1 | H | ASN | 76 | OD1-NH1 | C | ARG | 64 | 3.8 | H-bond |
| 1 | H | ASN | 76 | OD1-NE | C | ARG | 64 | 4.1 | H-bond |
| 1 | H | ARG | 94 | NH1-O | D | SER | 94 | 2.8 | H-bond |
| 1 | H | ARG | 94 | NH2-O | D | SER | 94 | 3.8 | H-bond |
| 1 | H | ASN | 97 | ND2-O | D | SER | 93 | 3.6 | H-bond |
| 1 | H | ASN | 97 | ND2-OG | D | SER | 95A | 4.4 | H-bond |
| 1 | L | SER | 56 | OG-OG | D | SER | 27 | 2.5 | H-bond |
| 2 | H | ARG | 19 | NH1-NH1 | O | ARG | 19 | 3.62 | Van-der-Waals |
| 2 | H | ARG | 19 | NH1-OE1 | O | GLN | 81 | 3.39 | H-bond |
| 2 | H | ARG | 19 | NH2-OE1 | O | GLN | 81 | 3.44 | H-bond |
| 2 | H | ASP | 72 | OD2-NE2 | O | HIS | 82A | 4.34 | H-bond |
| 2 | H | ARG | 75 | CD-CE1 | O | HIS | 82A | 3.41 | Van-der-Waals |
| 2 | H | TYR | 79 | OH-NE2 | O | GLN | 81 | 4.45 | H-bond |
| 2 | H | TYR | 79 | OH-CE1 | O | HIS | 82A | 4.32 | Van-der-Waals |
| 2 | H | GLN | 81 | OE1-NH1 | O | ARG | 19 | 3.38 | H-bond |
| 2 | H | GLN | 81 | OE1-NH2 | O | ARG | 19 | 3.43 | H-bond |
| 2 | H | GLN | 81 | NE2-OH | O | TYR | 79 | 4.44 | H-bond |
| 2 | H | HIS | 82A | NE2-OD2 | O | ASP | 72 | 4.32 | H-bond |
| 2 | H | HIS | 82A | CE1-CD | O | ARG | 75 | 3.4 | Van-der-Waals |
| 2 | H | HIS | 82A | CE1-OH | O | TYR | 79 | 4.31 | Van-der-Waals |
| 2 | H | GLN | 81 | NE2-OH | O | TYR | 79 | 4.44 | H-bond |
| 2 | H | HIS | 82A | NE2-OD2 | O | ASP | 72 | 4.32 | H-bond |
| 2 | H | HIS | 82A | CE1-CD | O | ARG | 75 | 3.4 | Van-der-Waals |
| 2 | H | HIS | 82A | CE1-OH | O | TYR | 79 | 4.31 | Van-der-Waals |

**Table S3-S9.** Complete set of homotypic contacts identified in both primary and secondary interfaces for all cryo-EM structures in this study.

**Table S4.** Homotypic contacts in 239-rsCSP cryo-EM structure

| Interface | Chain Fab 1 | Residue 1 | Position 1 | Atom | Chain Fab 2 | Residue 2 | Position 2 | Distance (Å) | Predicted interaction |
|---|---|---|---|---|---|---|---|---|---|
| **1** | H | GLN | 1 | O-O | Q | GLU | 64 | 4.0 | Van-der-Waals |
| **1** | H | GLN | 1 | N-O | Q | GLY | 65 | 3.0 | H-bond |
| **1** | H | VAL | 2 | CG2-CA | Q | GLY | 65 | 3.5 | Hydrophobic |
| **1** | H | VAL | 2 | N-O | Q | GLY | 65 | 4.1 | H-bond |
| **1** | H | VAL | 2 | CB-CA | Q | GLY | 65 | 4.4 | Hydrophobic |
| **1** | H | ARG | 26 | O-CA | Q | GLY | 65 | 3.5 | Van-der-Waals |
| **1** | H | ARG | 26 | O-N | Q | GLY | 65 | 4.1 | H-bond |
| **1** | H | LEU | 27 | CD1-OE1 | Q | GLU | 64 | 3.7 | Van-der-Waals |
| **1** | H | THR | 28 | CG2-O | Q | LYS | 57 | 3.8 | Van-der-Waals |
| **1** | H | THR | 28 | OG1-CE1 | Q | TYR | 59 | 3.5 | Van-der-Waals |
| **1** | H | THR | 28 | OG1-OE1 | Q | GLU | 64 | 3.5 | H-bond |
| **1** | H | THR | 28 | N-OE1 | Q | GLU | 64 | 3.8 | H-bond |
| **1** | H | ARG | 30 | NH2-O | Q | SER | 55 | 3.6 | H-bond |
| **1** | H | ARG | 30 | NE-O | Q | SER | 55 | 4.3 | H-bond |
| **1** | H | ARG | 30 | NE-OD1 | Q | ASN | 56 | 3.1 | H-bond |
| **1** | H | ARG | 30 | NH2-OD1 | Q | ASN | 56 | 4.2 | H-bond |
| **1** | H | ASN | 31 | ND2-O | Q | LYS | 57 | 2.3 | H-bond |
| **1** | H | ASN | 31 | ND2-CD1 | Q | PHE | 58 | 3.3 | Van-der-Waals |
| **1** | H | PHE | 32 | CE2-OE1 | Q | GLU | 64 | 3.1 | Van-der-Waals |
| **1** | H | PHE | 32 | CE1-NH2 | R | ARG | 95A | 4.1 | Van-der-Waals |
| **1** | H | LYS | 94 | NZ-CG | Q | GLU | 64 | 4.4 | Van-der-Waals |
| **1** | H | TRP | 96 | NE1-OD2 | Q | ASP | 61 | 3.0 | H-bond |
| **1** | H | TRP | 96 | CH2-N | R | ASP | 1 | 3.3 | Van-der-Waals |
| **1** | H | TRP | 96 | O-NH2 | R | ARG | 95A | 2.6 | H-bond |
| **1** | H | TRP | 96 | O-NH1 | R | ARG | 95A | 4.3 | H-bond |
| **1** | H | GLY | 97 | O-O | R | TYR | 94 | 4.2 | Van-der-Waals |
| **1** | H | GLY | 97 | CA-NH2 | R | ARG | 95A | 3.4 | Van-der-Waals |
| **1** | H | GLY | 97 | O-NH2 | R | ARG | 95A | 3.6 | H-bond |
| **1** | H | GLY | 97 | O-NH1 | R | ARG | 95A | 4.5 | H-bond |
| **1** | H | GLY | 98 | C-O | R | TYR | 94 | 4.4 | Van-der-Waals |
| **1** | H | GLY | 98 | N-NH2 | R | ARG | 95A | 4.2 | Van-der-Waals |
| **1** | H | ALA | 99 | CB-NE2 | R | GLN | 27 | 3.6 | Van-der-Waals |
| **1** | H | ALA | 99 | CA-O | R | TYR | 94 | 4.1 | Van-der-Waals |
| **1** | H | ALA | 99 | N-O | R | TYR | 94 | 4.3 | H-bond |
| **1** | H | ALA | 99 | CB-OG | R | SER | 95 | 3.3 | Van-der-Waals |
| **1** | H | ALA | 99 | N-OG | R | SER | 95 | 4.4 | H-bond |
| **1** | H | TYR | 102 | OH-O | Q | GLU | 64 | 4.0 | H-bond |
| **1** | L | ARG | 56 | NH1-O | Q | ASP | 61 | 3.1 | H-bond |
| **1** | L | ARG | 56 | NH2-O | Q | ASP | 61 | 3.8 | H-bond |
| **1** | L | ARG | 56 | NH1-CA | Q | SER | 62 | 3.2 | Van-der-Waals |
| **1** | L | ARG | 56 | NH1-O | Q | SER | 62 | 4.0 | H-bond |
| **1** | L | ARG | 56 | CD-NH2 | Q | ARG | 83 | 4.0 | Van-der-Waals |

**Table S5.** Homotypic contacts in 311-rsCSP cryo-EM structure

| Interface | Chain Fab 1 | Residue 1 | Position 1 | Atom | Chain Fab 2 | Residue 2 | Position 2 | Distance (Å) | Predicted interaction |
|---|---|---|---|---|---|---|---|---|---|
| 1 | H | GLY | 26 | O-O | Q | GLU | 64 | 4.2 | Van-der-Waals |
| 1 | H | GLY | 26 | O-CA | Q | GLY | 65 | 3.6 | Van-der-Waals |
| 1 | H | GLY | 26 | O-N | Q | GLY | 65 | 4.0 | H-bond |
| 1 | H | PHE | 27 | CB-OE2 | Q | GLU | 64 | 4.1 | Van-der-Waals |
| 1 | H | THR | 28 | OG1-CE1 | Q | TYR | 59 | 3.8 | Van-der-Waals |
| 1 | H | THR | 28 | OG1-OE2 | Q | GLU | 64 | 3.5 | H-bond |
| 1 | H | THR | 28 | N-OE2 | Q | GLU | 64 | 3.6 | H-bond |
| 1 | H | SER | 30 | O-NH2 | Q | ARG | 56 | 3.3 | H-bond |
| 1 | H | ASN | 31 | OD1-NH2 | Q | ARG | 56 | 3.1 | H-bond |
| 1 | H | ASN | 31 | OD1-NE | Q | ARG | 56 | 4.4 | H-bond |
| 1 | H | ASN | 31 | ND2-O | Q | ASN | 57 | 2.9 | H-bond |
| 1 | H | ASN | 31 | ND2-CD2 | Q | PHE | 58 | 3.7 | Van-der-Waals |
| 1 | H | TYR | 32 | OH-OE1 | Q | GLU | 64 | 3.2 | H-bond |
| 1 | H | TYR | 32 | OH-OE2 | Q | GLU | 64 | 3.4 | H-bond |
| 1 | H | TYR | 32 | OH-OG | R | SER | 95A | 4.4 | H-bond |
| 1 | H | TYR | 52A | CD1-NH2 | Q | ARG | 56 | 4.5 | Van-der-Waals |
| 1 | H | TYR | 97 | O-O | R | ARG | 94 | 4.2 | Van-der-Waals |
| 1 | H | TYR | 98 | O-CG | R | ARG | 94 | 3.3 | Van-der-Waals |
| 1 | H | TYR | 98 | OH-NH1 | R | ARG | 94 | 3.4 | H-bond |
| 1 | H | TYR | 98 | OH-NE | R | ARG | 94 | 4.1 | H-bond |
| 1 | H | TYR | 98 | OH-NH2 | R | ARG | 94 | 4.2 | H-bond |
| 1 | H | TYR | 98 | O-NE | R | ARG | 94 | 4.2 | H-bond |
| 1 | H | ASP | 99 | OD2-CB | R | SER | 27 | 4.4 | Van-der-Waals |
| 1 | H | ASP | 99 | OD2-NH2 | R | ARG | 93 | 3.0 | Salt-Bridge |
| 1 | H | ASP | 99 | OD2-NE | R | ARG | 93 | 4.3 | H-bond |
| 1 | H | ASP | 99 | OD1-NH2 | R | ARG | 94 | 2.5 | Salt-Bridge |
| 1 | H | ASP | 99 | OD1-NE | R | ARG | 94 | 3.0 | H-bond |
| 1 | H | ASP | 99 | OD2-NE | R | ARG | 94 | 3.2 | H-bond |
| 1 | H | ASP | 99 | OD2-NH2 | R | ARG | 94 | 3.6 | Salt-Bridge |
| 1 | H | ASP | 99 | OD1-NH1 | R | ARG | 94 | 4.2 | Salt-Bridge |
| 2 | H | ARG | 16 | NH2-O | X | THR | 18 | 3.5 | H-bond |
| 2 | H | ARG | 19 | NH2-CB | X | SER | 67 | 3.7 | Van-der-Waals |
| 2 | H | ARG | 19 | NH1-CB | X | SER | 65 | 4.1 | Van-der-Waals |
| 2 | H | ARG | 19 | NH2-OG | X | SER | 67 | 4.2 | H-bond |
| 2 | H | ARG | 19 | NH2-O | X | LYS | 66 | 4.3 | H-bond |
| 2 | H | ARG | 19 | NH1-OG | X | SER | 65 | 4.4 | H-bond |

**Table S6.** Homotypic contacts in 334-rsCSP cryo-EM structure

| Interface | Chain Fab 1 | Residue 1 | Position 1 | Atom | Chain Fab 2 | Residue 2 | Position 2 | Distance (Å) | Predicted interaction |
|---|---|---|---|---|---|---|---|---|---|
| **1** | H | ASN | 31 | OD1-OH | O | TYR | 58 | 2.6 | H-bond |
| **1** | H | ASN | 31 | ND2-CD | O | LYS | 56 | 4.2 | Van-der-Waals |
| **1** | H | ASN | 31 | OD1-NZ | O | LYS | 56 | 4.3 | H-bond |
| **1** | H | ASN | 31 | ND2-OH | O | TYR | 58 | 4.3 | H-bond |
| **1** | H | SER | 98 | O-CB | P | TYR | 94 | 3.8 | Van-der-Waals |
| **1** | H | SER | 99 | CB-OG | P | SER | 95 | 3.9 | Van-der-Waals |
| **1** | H | SER | 99 | CA-O | P | TYR | 94 | 4.1 | Van-der-Waals |
| **1** | H | SER | 99 | O-OE1 | P | GLN | 27 | 4.2 | Van-der-Waals |
| **1** | H | SER | 99 | O-OG | P | SER | 95 | 4.4 | H-bond |
| **1** | H | SER | 100 | O-CG | P | TYR | 94 | 3.6 | Van-der-Waals |
| **2** | H | ARG | 16 | NH1-OD1 | T | ASP | 17 | 3.1 | Salt-Bridge |
| **2** | H | ARG | 16 | CG-CD | T | ARG | 18 | 3.8 | Van-der-Waals |
| **2** | H | ARG | 16 | NH1-OD2 | T | ASP | 17 | 4.4 | Salt-Bridge |
| **2** | H | ARG | 16 | NH1-O | T | ARG | 18 | 4.5 | H-bond |
| **2** | H | SER | 17 | O-NE | T | ARG | 18 | 3.0 | H-bond |
| **2** | H | SER | 17 | OG-OG1 | T | THR | 20 | 3.7 | H-bond |
| **2** | H | SER | 17 | O-NH2 | T | ARG | 18 | 3.8 | H-bond |
| **2** | H | LEU | 18 | CD1-NH2 | T | ARG | 18 | 3.4 | Van-der-Waals |
| **2** | H | THR | 19 | OG1-OG | T | SER | 65 | 3.2 | H-bond |
| **2** | H | THR | 19 | CG2-OG1 | T | THR | 74 | 4.1 | Van-der-Waals |
| **2** | H | THR | 68 | CG2-CG | T | GLU | 70 | 3.8 | Van-der-Waals |
| **2** | H | SER | 70 | CB-O | T | SER | 67 | 3.5 | Van-der-Waals |
| **2** | H | SER | 70 | OG-N | T | SER | 67 | 3.5 | H-bond |
| **2** | H | SER | 70 | OG-CA | T | GLY | 66 | 3.8 | Van-der-Waals |
| **2** | H | SER | 70 | OG-O | T | SER | 67 | 3.9 | H-bond |
| **2** | H | ARG | 71 | O-OG | T | SER | 67 | 3.2 | H-bond |
| **2** | H | ASP | 72 | OD2-CG2 | T | THR | 31 | 3.2 | Van-der-Waals |
| **2** | H | LYS | 75 | CE-CB | T | SER | 52 | 3.9 | Van-der-Waals |
| **2** | H | TYR | 79 | CD2-CB | T | SER | 65 | 3.7 | Van-der-Waals |
| **2** | H | TYR | 79 | OH-OG | T | SER | 52 | 3.9 | H-bond |
| **2** | H | TYR | 79 | CG-N | T | GLY | 66 | 4.3 | Van-der-Waals |
| **2** | H | GLN | 81 | NE2-CA | T | GLY | 66 | 3.0 | Van-der-Waals |
| **2** | H | GLN | 81 | NE2-C | T | SER | 65 | 3.6 | Van-der-Waals |
| **2** | H | GLN | 81 | NE2-O | T | SER | 65 | 3.7 | H-bond |
| **2** | H | GLN | 81 | NE2-OG | T | SER | 65 | 3.7 | H-bond |
| **2** | H | GLN | 81 | CG-OG1 | T | THR | 72 | 3.9 | Van-der-Waals |
| **2** | H | GLN | 81 | OE1-CG | T | GLU | 70 | 4.1 | Van-der-Waals |
| **2** | H | GLN | 81 | NE2-O | T | GLU | 70 | 4.4 | H-bond |

**Table S7.** Homotypic contacts in 337-rsCSP cryo-EM structure

| Interface | Chain Fab 1 | Residue 1 | Position 1 | Atom | Chain Fab 2 | Residue 2 | Position 2 | Distance (Å) | Predicted interaction |
|---|---|---|---|---|---|---|---|---|---|
| **1** | H | SER | 28 | OG-N | M | LYS | 57 | 3.3 | H-bond |
| **1** | H | SER | 28 | OG-O | M | SER | 55 | 3.7 | H-bond |
| **1** | H | SER | 28 | OG-O | M | LYS | 57 | 3.8 | H-bond |
| **1** | H | SER | 28 | OG-CA | M | LYS | 56 | 3.9 | Van-der-Waals |
| **1** | H | SER | 30 | OG-CG | M | LYS | 56 | 3.3 | Van-der-Waals |
| **1** | H | SER | 30 | OG-O | M | SER | 55 | 3.3 | H-bond |
| **1** | H | SER | 30 | OG-N | M | LYS | 56 | 4.0 | H-bond |
| **1** | H | SER | 30 | OG-OG | M | SER | 55 | 4.4 | H-bond |
| **1** | H | THR | 31 | CG2-CG | M | LYS | 56 | 3.9 | Van-der-Waals |
| **1** | H | THR | 31 | CG2-CZ | M | PHE | 58 | 4.1 | Van-der-Waals |
| **1** | H | THR | 31 | CG2-O | M | LYS | 57 | 4.1 | Van-der-Waals |
| **1** | H | TYR | 32 | OH-NH2 | M | ARG | 64 | 2.5 | H-bond |
| **1** | H | TYR | 32 | OH-NH1 | M | ARG | 64 | 4.3 | H-bond |
| **1** | H | HIS | 52 | CD2-CE | M | LYS | 56 | 4.3 | Van-der-Waals |
| **1** | H | TYR | 99 | OH-NH2 | M | ARG | 64 | 3.8 | H-bond |
| **1** | H | TYR | 99 | CZ-CE2 | M | PHE | 58 | 4.0 | Hydrophobic |
| **1** | H | TYR | 99 | CE1-CD2 | M | PHE | 58 | 4.0 | Hydrophobic |
| **1** | H | TYR | 99 | CE1-CE2 | M | PHE | 58 | 4.1 | Hydrophobic |
| **1** | H | TYR | 99 | CE2-CE2 | M | PHE | 58 | 4.2 | Hydrophobic |
| **1** | H | TYR | 99 | CZ-CD2 | M | PHE | 58 | 4.2 | Hydrophobic |
| **1** | H | TYR | 99 | CD1-CD2 | M | PHE | 58 | 4.4 | Hydrophobic |
| **1** | H | TYR | 99 | CD1-CE2 | M | PHE | 58 | 4.4 | Hydrophobic |
| **1** | H | TYR | 99 | CD1-CZ2 | N | TRP | 94 | 3.8 | Hydrophobic |
| **1** | H | TYR | 99 | CD1-CH2 | N | TRP | 94 | 3.9 | Hydrophobic |
| **1** | H | TYR | 99 | CE1-CH2 | N | TRP | 94 | 4.3 | Hydrophobic |
| **1** | H | PHE | 100 | CZ-O | N | THR | 93 | 2.8 | Van-der-Waals |
| **1** | H | PHE | 100 | CE1-NE1 | N | TRP | 94 | 3.2 | Van-der-Waals |
| **1** | H | PHE | 100 | CE1-CD1 | N | TRP | 94 | 3.6 | Hydrophobic |
| **1** | H | PHE | 100 | CE1-CE2 | N | TRP | 94 | 3.8 | Hydrophobic |
| **1** | H | PHE | 100 | CD1-CD1 | N | TRP | 94 | 4.2 | Hydrophobic |
| **1** | H | PHE | 100 | CE1-CG | N | TRP | 94 | 4.4 | Hydrophobic |
| **1** | H | PHE | 100 | CE1-CZ2 | N | TRP | 94 | 4.4 | Hydrophobic |
| **1** | H | PHE | 100 | CZ-CD1 | N | TRP | 94 | 4.5 | Hydrophobic |
| **2** | H | ARG | 16 | NH1-NH1 | R | ARG | 18 | 2.9 | Van-der-Waals |
| **2** | H | ARG | 16 | NH1-CG2 | R | THR | 74 | 3.6 | Van-der-Waals |
| **2** | H | ARG | 16 | NH1-OG1 | R | THR | 74 | 4.5 | H-bond |
| **2** | H | SER | 17 | CB-OG | R | SER | 65 | 3.9 | Van-der-Waals |
| **2** | H | ARG | 19 | NE-OG | R | SER | 67 | 2.8 | H-bond |
| **2** | H | ARG | 19 | NH2-OG | R | SER | 31 | 2.8 | H-bond |
| **2** | H | ARG | 19 | NE-OG | R | SER | 31 | 3.5 | H-bond |
| **2** | H | ARG | 19 | NH1-OG | R | SER | 31 | 4.1 | H-bond |
| **2** | H | ARG | 19 | NH2-OG | R | SER | 67 | 4.4 | H-bond |
| **2** | H | ARG | 19 | CD-O | R | GLY | 66 | 4.5 | Van-der-Waals |
| **2** | H | GLN | 81 | NE2-CA | R | GLY | 66 | 3.1 | Van-der-Waals |
| **2** | H | GLN | 81 | NE2-N | R | SER | 67 | 3.3 | Van-der-Waals |
| **2** | H | GLN | 81 | OE1-OG | R | SER | 67 | 3.9 | H-bond |
| **2** | H | GLN | 81 | OE1-N | R | SER | 67 | 4.1 | H-bond |
| **2** | H | GLN | 81 | NE2-O | R | GLY | 66 | 4.2 | H-bond |

**Table S8.** Homotypic contacts in 356-rsCSP cryo-EM structure

| Interface | Chain Fab 1 | Residue 1 | Position 1 | Atom | Chain Fab 2 | Residue 2 | Position 2 | Distance (Å) | Predicted interaction |
|---|---|---|---|---|---|---|---|---|---|
| 1 | H | GLY | 26 | O-O | Q | GLU | 64 | 4.2 | Van-der-Waals |
| 1 | H | GLY | 26 | O-CA | Q | GLY | 65 | 3.6 | Van-der-Waals |
| 1 | H | GLY | 26 | O-N | Q | GLY | 65 | 4.3 | H-bond |
| 1 | H | PHE | 27 | CB-CG | Q | GLU | 64 | 4.0 | Van-der-Waals |
| 1 | H | PHE | 27 | CA-CA | Q | GLY | 65 | 4.3 | Hydrophobic |
| 1 | H | THR | 28 | CG2-O | Q | LYS | 57 | 4.4 | Van-der-Waals |
| 1 | H | THR | 28 | OG1-CE1 | Q | TYR | 59 | 3.2 | Van-der-Waals |
| 1 | H | THR | 28 | OG1-OE2 | Q | GLU | 64 | 2.9 | H-bond |
| 1 | H | THR | 28 | N-OE2 | Q | GLU | 64 | 3.4 | H-bond |
| 1 | H | ARG | 30 | NH1-O | Q | SER | 55 | 3.6 | H-bond |
| 1 | H | ARG | 30 | NH2-O | Q | SER | 55 | 4.2 | H-bond |
| 1 | H | ARG | 30 | NH1-CA | Q | ASN | 56 | 4.1 | Van-der-Waals |
| 1 | H | ARG | 30 | NH1-OD1 | Q | ASN | 56 | 4.2 | H-bond |
| 1 | H | ARG | 30 | NH2-OD1 | Q | ASN | 56 | 4.3 | H-bond |
| 1 | H | ASN | 31 | ND2-O | Q | LYS | 57 | 2.7 | H-bond |
| 1 | H | ASN | 31 | OD1-CE2 | Q | PHE | 58 | 4.2 | Van-der-Waals |
| 1 | H | PHE | 32 | CZ-OE1 | Q | GLU | 64 | 3.8 | Van-der-Waals |
| 1 | H | LEU | 97 | O-OD2 | Q | ASP | 61 | 3.8 | Van-der-Waals |
| 1 | H | PHE | 98 | CB-CB | Q | ASP | 61 | 3.7 | Van-der-Waals |
| 1 | H | PHE | 98 | CE1-N | R | GLN | 1 | 3.8 | Van-der-Waals |
| 1 | H | TYR | 99 | O-NE2 | R | GLN | 1 | 3.1 | H-bond |
| 1 | H | TYR | 99 | CE2-OD1 | R | ASN | 93 | 4.0 | Van-der-Waals |
| 1 | H | ASP | 100 | OD1-CA | R | GLN | 1 | 3.4 | Van-der-Waals |
| 1 | H | ASP | 100 | OD1-N | R | GLN | 1 | 4.2 | H-bond |
| 1 | H | ASP | 100 | OD2-NH2 | R | ARG | 27 | 2.7 | Salt-Bridge |
| 1 | H | ASP | 100 | OD1-NH1 | R | ARG | 27 | 2.9 | Salt-Bridge |
| 1 | H | ASP | 100 | OD1-NH2 | R | ARG | 27 | 3.3 | Salt-Bridge |
| 1 | H | ASP | 100 | OD2-NH1 | R | ARG | 27 | 3.6 | Salt-Bridge |
| 1 | H | ASP | 100 | CB-NH2 | R | ARG | 27 | 3.6 | Van-der-Waals |
| 1 | H | ASP | 100 | OD2-ND2 | R | ASN | 93 | 3.8 | H-bond |
| 1 | H | ASP | 100 | OD1-ND2 | R | ASN | 93 | 4.1 | H-bond |
| 1 | H | HIS | 100A | N-ND2 | R | ASN | 93 | 3.6 | Van-der-Waals |
| 1 | H | HIS | 100A | N-OD1 | R | ASN | 93 | 3.6 | H-bond |
| 1 | H | ASP | 100B | OD2-NH2 | R | ARG | 27 | 3.9 | Salt-Bridge |
| 1 | H | ASP | 100B | N-ND2 | R | ASN | 93 | 4.4 | Van-der-Waals |
| 2 | H | GLN | 13 | NE2-OE2 | X | GLU | 17 | 3.5 | H-bond |
| 2 | H | ARG | 16 | NH1-CG | X | GLU | 17 | 4.0 | Van-der-Waals |
| 2 | H | ARG | 16 | NH1-OE1 | X | GLU | 17 | 4.1 | Salt-Bridge |
| 2 | H | ARG | 16 | NH1-O | X | ARG | 18 | 4.1 | H-bond |
| 2 | H | SER | 17 | O-CD | X | ARG | 18 | 4.5 | Van-der-Waals |
| 2 | H | SER | 17 | OG-CG2 | X | THR | 20 | 3.2 | Van-der-Waals |
| 2 | H | SER | 17 | OG-OG1 | X | THR | 20 | 3.3 | H-bond |
| 2 | H | SER | 17 | O-OG1 | X | THR | 20 | 4.2 | H-bond |
| 2 | H | LEU | 18 | CD1-NH2 | X | ARG | 18 | 4.3 | Van-der-Waals |
| 2 | H | ARG | 19 | NE-OG | X | SER | 65 | 3.1 | H-bond |
| 2 | H | ARG | 19 | NH1-OG | X | SER | 65 | 3.6 | H-bond |
| 2 | H | ARG | 19 | NH2-OG | X | SER | 65 | 3.9 | H-bond |
| 2 | H | ARG | 19 | NH2-N | X | GLY | 66 | 3.6 | Van-der-Waals |
| 2 | H | ARG | 19 | NH1-OG1 | X | THR | 72 | 4.2 | H-bond |
| 2 | H | LYS | 75 | NZ-OG | X | SER | 52 | 4.3 | H-bond |
| 2 | H | GLN | 81 | OE1-OG1 | X | THR | 72 | 4.1 | H-bond |

**Table S9.** Homotypic contacts in 364-rsCSP cryo-EM structure

| Interface | Chain Fab 1 | Residue 1 | Position 1 | Atom | Chain Fab 2 | Residue 2 | Position 2 | Distance (Å) | Predicted interaction |
|---|---|---|---|---|---|---|---|---|---|
| **1** | H | PHE | 27 | O-OD2 | E | ASP | 61 | 3.4 | Van-der-Waals |
| **1** | H | PHE | 27 | N-OD2 | E | ASP | 61 | 4.3 | H-bond |
| **1** | H | PHE | 27 | C-OD2 | F | ASP | 1 | 4.0 | Van-der-Waals |
| **1** | H | THR | 28 | CA-OD1 | E | ASP | 61 | 3.4 | Van-der-Waals |
| **1** | H | THR | 28 | OG1-OD1 | E | ASP | 61 | 3.5 | H-bond |
| **1** | H | THR | 28 | N-OD2 | E | ASP | 61 | 3.9 | H-bond |
| **1** | H | THR | 28 | N-OD1 | E | ASP | 61 | 4.0 | H-bond |
| **1** | H | THR | 28 | OG1-OD2 | E | ASP | 61 | 4.3 | H-bond |
| **1** | H | THR | 28 | CG2-O | E | TYR | 59 | 4.5 | Van-der-Waals |
| **1** | H | THR | 28 | OG1-OD2 | F | ASP | 1 | 2.4 | H-bond |
| **1** | H | THR | 28 | OG1-OD1 | F | ASP | 1 | 3.0 | H-bond |
| **1** | H | THR | 28 | N-OD2 | F | ASP | 1 | 3.2 | H-bond |
| **1** | H | THR | 28 | N-OD1 | F | ASP | 1 | 3.5 | H-bond |
| **1** | H | THR | 28 | CG2-CZ | F | TYR | 94 | 3.6 | Van-der-Waals |
| **1** | H | SER | 30 | OG-NE2 | E | GLN | 64 | 3.3 | H-bond |
| **1** | H | SER | 30 | OG-OE1 | E | GLN | 64 | 4.4 | H-bond |
| **1** | H | GLY | 31 | CA-OH | F | TYR | 94 | 3.4 | Van-der-Waals |
| **1** | H | GLY | 31 | O-NH2 | F | ARG | 93 | 3.5 | H-bond |
| **1** | H | GLY | 31 | N-OH | F | TYR | 94 | 4.1 | H-bond |
| **1** | H | GLY | 31 | O-OH | F | TYR | 94 | 4.3 | H-bond |
| **1** | H | GLY | 31 | CA-CZ | F | TYR | 94 | 4.4 | Hydrophobic |
| **1** | H | TYR | 32 | OH-NE | F | ARG | 93 | 3.6 | H-bond |
| **1** | H | TYR | 32 | OH-NH2 | F | ARG | 93 | 3.6 | H-bond |
| **1** | H | TYR | 32 | CE1-OD1 | F | ASP | 1 | 3.8 | Van-der-Waals |
| **1** | H | TYR | 32 | OH-OD1 | F | ASP | 1 | 3.9 | H-bond |
| **1** | H | TYR | 32 | OH-N | F | ASP | 1 | 4.1 | H-bond |
| **1** | H | ASN | 73 | O-CG | E | GLN | 64 | 4.0 | Van-der-Waals |
| **1** | H | LYS | 76 | NZ-CG | E | ASP | 61 | 4.2 | Van-der-Waals |
| **1** | H | LYS | 76 | NZ-OD2 | E | ASP | 61 | 4.3 | Salt-Bridge |
| **1** | H | ASP | 97 | OD2-OE1 | F | GLN | 27 | 2.7 | Van-der-Waals |
| **1** | H | ASP | 97 | OD2-NE2 | F | GLN | 27 | 3.3 | H-bond |
| **1** | H | ASP | 97 | OD1-NE | F | ARG | 93 | 4.1 | H-bond |

| Table S10. Summary of biolayer interferometry binding and liver burden data | | | | | | | | | | |
|---|---|---|---|---|---|---|---|---|---|---|
| | $NPNA_4$ | | | $NPNA_8$ | | | rsCSP | | | |
| mAb | $k_{ON}$ (1/Ms) | $k_{OFF}$ (1/s) | $K_D$ (nM) | $k_{ON}$ (1/Ms) | $k_{OFF}$ (1/s) | $K_D$ (nM) | $k_{ON}$ (1/Ms) | $k_{OFF}$ (1/s) | $K_D$ (nM) | Liver burden (% inh.) |
| 227 | 1.3E+05 | 5.8E-03 | 46.3 | 4.0E+04 | 3.5E-03 | 80.3 | 1.7E+05 | 2.5E-04 | 1.6 | 52.5 |
| 239 | 1.1E+05 | 6.3E-03 | 44.7 | 8.3E+04 | 5.6E-04 | 7.2 | 1.4E+05 | 6.3E-05 | 0.5 | 81 |
| 239R | 4.1E+05 | 3.3E-02 | 165 | 1.2E+05 | 2.2E-02 | 205.0 | 3.3E+05 | 2.9E-02 | 110.0 | 31.2 |
| 311 | 2.7E+04 | 4.1E-03 | 212 | 6.6E+04 | 2.5E-04 | 6.1 | 6.6E+04 | 7.6E-05 | 1.7 | 87.7 |
| 311R | 7.1E+04 | 8.8E-03 | 131 | 7.1E+04 | 5.1E-03 | 401.0 | 6.5E+04 | 5.6E+03 | 99.0 | 41.4 |
| 334 | 5.3E+04 | 2.1E-03 | 49.9 | 5.2E+04 | 1.3E-03 | 22.4 | 1.3E+05 | 1.1E-03 | 4.9 | 83.6 |
| 337 | 1.3E+05 | 7.0E-03 | 36.9 | 6.6E+04 | 3.5E-03 | 39.4 | 1.5E+05 | 1.3E-03 | 5.6 | 68.4 |
| 356 | 8.0E+04 | 1.3E-02 | 123 | 8.7E+04 | 5.6E-04 | 6.4 | 1.4E+05 | 1.1E-04 | 0.8 | 87.7 |
| 356R | 5.6E+05 | 3.0E-02 | 65 | 7.9E+04 | 1.7E-02 | 124.0 | 1.6E+05 | 1.6E-02 | 67.0 | 55.9 |
| 364 | 4.3E+04 | 2.7E-03 | 260 | 6.5E+04 | 2.7E-04 | 4.9 | 1.3E+05 | 1.2E-04 | 0.9 | 85.4 |
| 317 | 8.5E+04 | 3.0E-04 | 3.8 | 6.9E+04 | 1.9E-04 | 2.9 | 8.3E+05 | 3.4E-04 | 0.4 | 91.4 |

**Table S10.** Summary of biolayer interferometry binding and liver burden data. Kinetic parameters are averages across at least 4 fits with $R^2 \geq 0.98$.

**Table S11. X-ray data collection and refinement statistics (molecular replacement) for 311R Fab**

| | 311R-(NPNA)3 |
|---|---|
| **Data collection** | |
| Space group | $P2_12_12_1$ |
| Cell dimensions | |
| *a*, *b*, *c* (Å) | 45.41, 70.77, 170.25 |
| α, β, γ (°) | 90.00, 90.00, 90.00 |
| Resolution (Å) | 1.90(1.94-1.90)* |
| $R_{sym}$ or $R_{merge}$ | 0.055 (0.625) |
| $I / \sigma I$ | 17.2(1.6) |
| Completeness (%) | 99.8(97.6) |
| Redundancy | 3.3(2.3) |
| **Refinement** | |
| Resolution (Å) | 1.90 |
| No. reflections | 44216 |
| $R_{work}$ / $R_{free}$ | 0.2199 / 0.2534 |
| No. atoms | 3312 |
| Protein | 3312 |
| Ligand/ion | |
| Water | |
| *B*-factors | (Ask for input) |
| Protein | |
| Ligand/ion | |
| Water | |
| R.m.s. deviations | |
| Bond lengths (Å) | 0.0108 |
| Bond angles (°) | 1.7354 |

*Single crystal. *Values in parentheses are for highest-resolution shell.

[AU: Equations defining various *R*-values are standard and hence are no longer defined in the footnotes.]
[AU: Ramachandran statistics should be in Methods section at the end of Refinement subsection.]
[AU: Wavelength of data collection, temperature and beamline should all be in Methods section.]
